# SpliceWiz: easy, optimized, and accurate alternative splicing analysis in R

**DOI:** 10.1101/2022.07.05.498887

**Authors:** Alex CH Wong, Justin J-L Wong, John EJ Rasko, Ulf Schmitz

## Abstract

Despite an abundance of publicly available RNA sequencing datasets, a lack of integrated user-friendly tools hinder exploration of alternative splicing. *SpliceWiz* is an innovative, ultra-fast graphical R application that accurately quantifies splicing events using isoform-specific alignments. It is designed to accommodate hundreds of samples typically seen in clinical datasets. Novel event filters remove low-confidence measurements from analysis, enhancing accuracy over existing methods. Group-averaged strand-specific sequencing coverage plots enable clear visualization of group differences in alternative splicing, using a new file format with demonstrable performance improvements over the current BigWig standard. *ompBAM*, a C++ library upon which *SpliceWiz* is built, automates multi-threaded alignment file processing for R package developers. *SpliceWiz* is a powerful platform for diverse users to explore alternative splicing in large datasets.

## INTRODUCTION

Alternative splicing (AS) is a highly regulated process that diversifies the transcriptome, allowing a single gene to express multiple protein-coding and non-coding RNA isoforms. Splicing is performed and regulated by the spliceosome, a megadalton complex of ribonucleoproteins and splicing protein co-factors [1]. Splice sites demarcating exon-intron boundaries are recognized, leading to the excision of introns and selection of exons that remain in messenger RNA (mRNA) transcripts [2]. AS arises when splice sites are differentially recognized, resulting in alternate combinations of exons and thereby different mRNA transcript isoforms. AS regulates multiple critical biological processes including cellular development and differentiation [3], apoptosis [4], adaptation to physiological stress [5], and reproduction [6]. Intron retention (IR) is a special case of AS where one or more introns remain as part of mature mRNA transcripts. In contrast to other forms of AS, IR leads to the finetuning of gene expression through mechanisms including sequestration and/or degradation of mRNAs [7, 8]. There is an ever-increasing interest in robust quantitation of AS and IR levels in biological samples, including meaningfully large datasets. Underpinning this interest is the development of computationally efficient, accurate, user-friendly and well-maintained bioinformatic tools.

At its most basic form, alternative splicing events (ASEs) are conceptualized as binary choices, of which there are seven main forms [9] (Figure S1). Five of these depend on splice site choice alone; comprising skipped exons (SE), mutually exclusive exons (MXE), alternative 5’- and 3’- splice site usage (A5SS / A3SS), and IR. Additionally, alternative first exon (AFE) events arise from differential transcriptional start site usage followed by splicing, whereas alternate last exon (ALE) events arise when differential 3’-splice site usage triggers alternative polyadenylation. Apart from IR, all the above ASEs can be quantified using counts of reads aligned across splice junctions (hereafter junction reads) in short-read RNA sequencing (RNA-seq) [10–12]. The advantage of this approach is that junction reads can be unambiguously assigned to a specific splicing event. This approach contrasts with transcript quantitation, where most reads cannot be assigned definitively to any one transcript, and their accuracy is limited by the assumptions of the probabilistic models employed as well as the complexity of transcript annotations.

Currently available bioinformatics tools have several disadvantages that limit their use, especially when analysing large datasets. These include (A) computationally inefficient processes due to their implementation in high-level programming languages and lack of multi-threading; (B) inability to perform statistical analysis using complex experimental designs; (C) rigid and non-interactive user interfaces limiting explorative data analysis; (D) inability to visualize differential ASEs as aggregate changes in RNA-seq coverage across biological groups or experimental replicates, and (E) a steep software learning curve due to lack of a graphical user interface (GUI). Most *R* packages for AS rely on the *RSamtools* package [13] and the *Rhtslib* C library [14] to access Binary Alignment Map (BAM) files. These do not currently offer multi-threaded processing. Moreover, tools that specialize in measuring IR, including *IRFinder* [15], *IntEREst* [16], and *iREAD* [17] do not concurrently quantify other forms of AS.

To address the above limitations, we created *SpliceWiz*, an accessible, user-friendly, interactive, and computationally efficient *R* package. *SpliceWiz* leverages *IRFinder* core concepts to quantify IR [15], and quantifies other forms of ASE using junction reads. It utilizes generalized linear model-based differential splicing analysis to cater for complex experimental designs. Counts are modeled using negative binomial, log-normal, or beta-binomial distributions via well-established statistical frameworks *DESeq2* [18], *limma* [19] and *DoubleExpSeq* [20], respectively. Novel filtering approaches are implemented to exclude low-confidence ASEs in complex transcriptomes, thereby enhancing accuracy. Additionally, novel group-averaged RNA-seq coverage plots visualize differential splicing across biological conditions. *SpliceWiz* is ultra-fast, employing multi-threaded processing of BAM files and a novel file format optimized to store strand-specific RNA-seq coverage data. It offers both a graphical user and command line interface, thereby reducing the learning curve for all users. Additionally, we provide *ompBAM*, a C++ application programming interface (API) on which *SpliceWiz* is built. *ompBAM* is an R package development tool that automates multi-threaded BAM processing. Overall, *SpliceWiz* is a computationally powerful and user-friendly AS and IR analysis and visualization tool optimized for large datasets.

## MATERIAL AND METHODS

### SpliceWiz implementation and workflow

The *SpliceWiz Suite* comprises the *SpliceWiz* R package and its companion C++ API *ompBAM* (Figure S2A). The *SpliceWiz* GUI provides point-and-click interactivity including searchable Ensembl references (Figure S2B), experiment construction using spreadsheet-like tables (Figure S2C), and inter-connected visualization features for data exploration (Figure S2D-F). *SpliceWiz* also provides an R-based command-line interface with equivalent functionality allowing for script-based automation. *ompBAM*, upon which *SpliceWiz* is built, manages the computational steps required for multi-threaded BAM file processing (Supplementary Methods). With *ompBAM*, developers can create R packages (within the Rcpp environment [21]) with the performance gains as seen in *SpliceWiz*.

The *SpliceWiz* analysis workflow is illustrated in Figure 1. Briefly, *SpliceWiz’s* splicing reference is generated from the genome and gene annotation files sourced either from the *Ensembl* FTP site or via *Bioconductor’s AnnotationHub* [22]. After read alignment using an external splice-aware genome aligner such as *STAR* [23], Rsubread [24] or *HISAT2* [25], BAM files are processed to quantify junction reads and intronic abundance estimation, compile quality control (QC) metrics, and record per-nucleotide alignment coverage. Results of individual samples are then collated into a H5 database to facilitate memory efficiency. Data import is managed using *NxtSE*, a specialized *SummarizedExperiment* S4 object tailored to store isoform-specific counts. Users can annotate samples to their experimental groups using the *NxtSE* object. ASE filtering is performed to remove low-confidence ASEs prior to differential analysis. Results of differential analysis and PSI values can then be visualized via scatter plots, volcano plots and heatmaps. Additionally, ASEs can be visualized using novel group-averaged coverage plots. The GUI enables further interactivity by linking ASEs of interest between these visualization modalities and provides coverage plots that are scrollable across the genomic axis.

**Figure 1:**
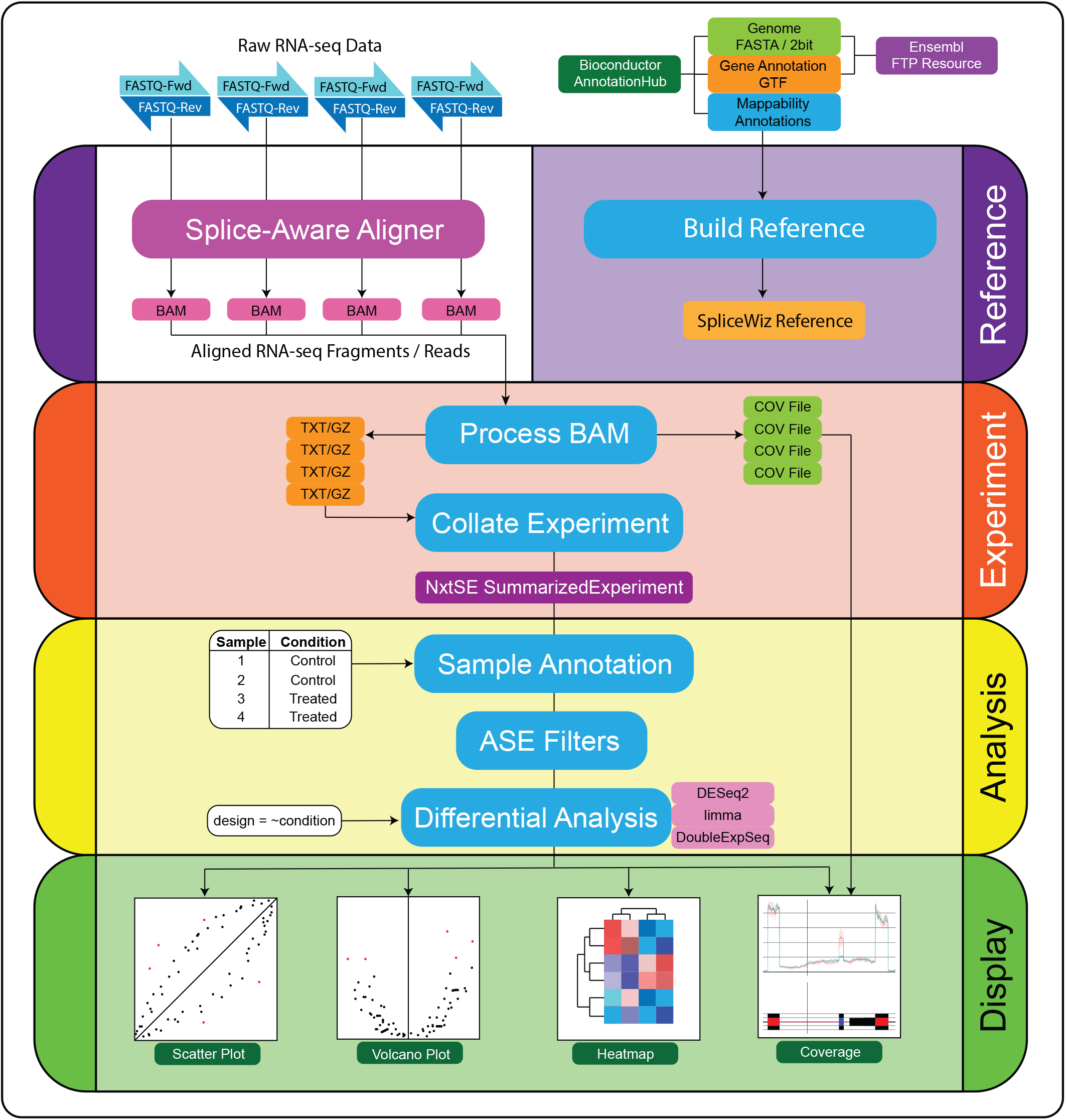
Conceptual overview of the *SpliceWiz* architecture and workflow. The *SpliceWiz* workflow comprises the construction of the *SpliceWiz* reference (Reference), processing of RNA-seq samples and collation of the experiment (Experiment), annotation and filtering of the *NxtSE* object prior to differential analysis (Analysis), and visualization of results (Display).

### Filter design to remove problematic alternative splicing events

*SpliceWiz* quantifies ASEs using the percent-spliced-in (PSI) metric (Supplementary Methods). PSI estimation in low-abundance genes or in samples sequenced at low depth typically exhibit higher measurement uncertainty (Figure 2A). In complex transcriptomes, two or more pairs of binary ASEs often overlap across the genomic axis. For example, alternative splice site usage may occur in exons flanking an alternatively skipped exon (Figure 2B). In such a scenario, one isoform is highly expressed (the major isoform), whereas the other two alternative isoforms are less abundant (minor isoforms). Analogous to low depth, the PSI estimates of ASE comprising two minor isoforms will also exhibit higher uncertainty. We define ASEs of two minor isoforms to be of low *participation* (Figure 2B). ASEs can be filtered using the *participation ratio* (Supplementary Methods).

**Figure 2:**
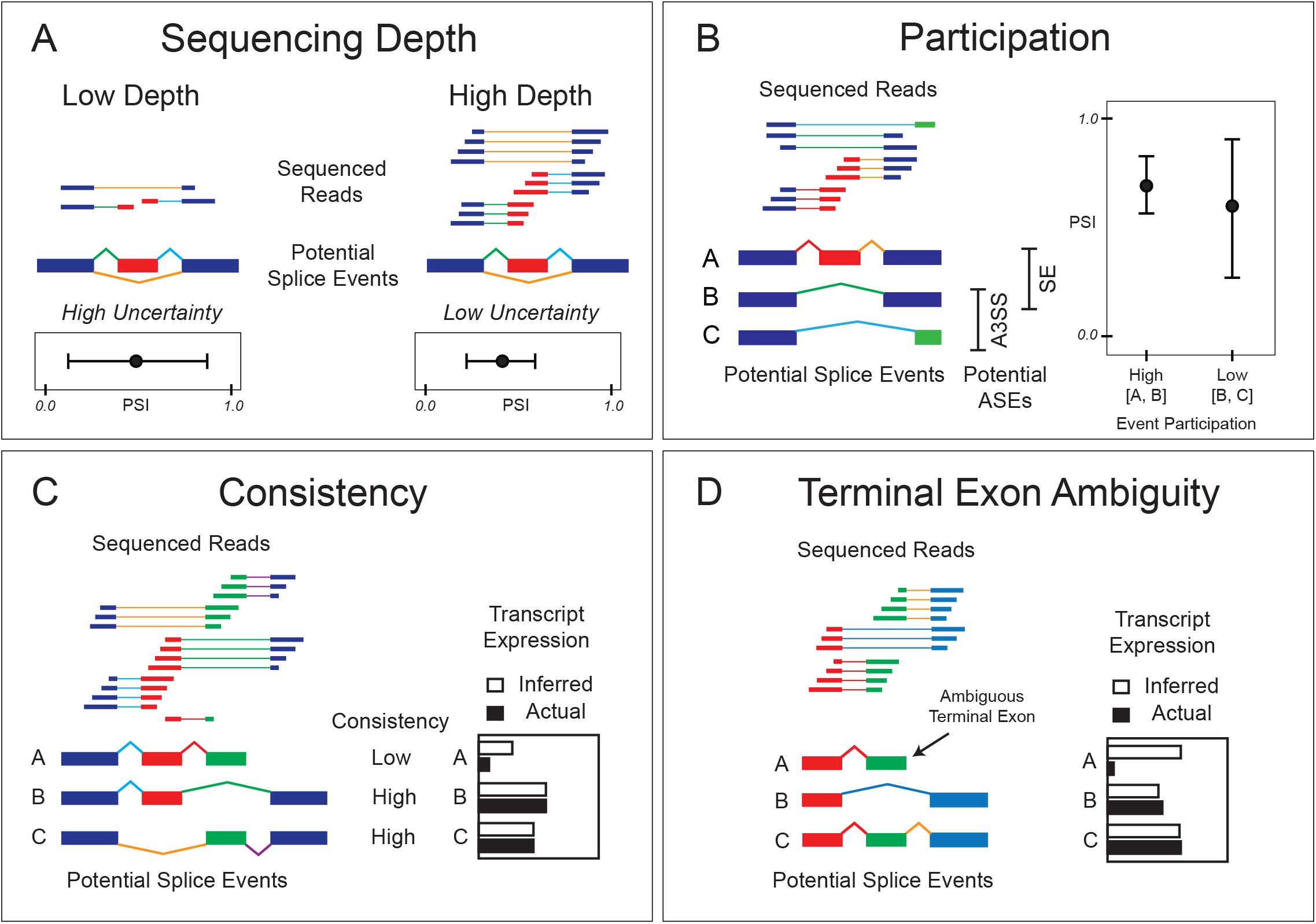
Factors that confound accuracy of splicing quantitation. When estimating isoform ratios using junction reads, measurement uncertainty of percent-spliced-in (PSI) increases when there is **(A)** low sequencing depth at the junctions of interest. Likewise, **(B)** when a pair of isoforms represents a low proportion of junction reads where multiple participating alternate splicing event (ASE) pairs exist, their combined low depth leads to measurement uncertainty. **(C)** Where the expression of an isoform relies on averaging counts of its tandem junctions, low consistency between these counts indicates confounding expression of isoforms that overlap one but not both tandem junctions. **(D)** Similarly, terminal exon expression is confounded when such exons are also the middle exons of other transcripts. Error bars indicate measurement uncertainty of PSI.

In skipped exons and mutually exclusive exons, isoform abundance is calculated by taking the average junction counts of the two consecutive introns (hereafter referred to as tandem junctions). If the isoform comprising these tandem junctions is dominantly expressed, it is expected that the read counts of both junctions are of similar magnitude. This comparison is quantified using the *consistency ratio* (Supplementary Methods). Isoforms with low *consistency ratio* typically arise when an alternate isoform, sharing one but not both introns with the isoform of interest, is expressed at levels that interfere with quantitation. Low-consistency isoforms cannot be reliably quantified from tandem junction counts alone (Figure 2C), and ASEs comprising low consistency isoforms can be identified and filtered using the *consistency ratio*.

Similarly, first and last introns of transcripts sometimes overlap with middle introns of other transcripts. Consequently, junction reads arising from these introns cannot be confidently attributed to expression of the corresponding first or last exons (Figure 2D). Using the gene annotations, AFE and ALE events comprising introns that overlap middle introns of other transcripts can be excluded from analysis (terminal exon filter).

For IR, *SpliceWiz* utilizes two filtering approaches analogous to participation and consistency for other ASE modalities. IR-participation is defined as the proportion of the measured intron that is covered by reads. IR-consistency is the consistency ratio between upstream and downstream counts that span across the donor and acceptor splice sites of the intron of interest (exon-intron spanning read counts). These two filters for IR complement those of other ASEs.

### COV file implementation for storing RNA-seq coverage data

The current implementation of *BigWig* (43) is not optimized for stranded RNA-seq coverage data, which requires the handling of positive, negative and unstranded coverage values on the same genomic axis. To enable streamlined read/write access of stranded coverage data, we designed the *COV* file format (Figure S3, Table S1). *COV* is a binary indexed file format utilizing *BGZF*-based compression, allowing random read access as in *BigWig*. However, *COV* accommodates 3 vectors of coverage data for positive, negative and unstranded RNA-seq coverage. Every *BGZF* block (64 kb) of compressed coverage data is indexed by the starting genomic coordinates of the data it contains and its file offsets, allowing for fast random-access data retrieval. *COV* files are generated concurrently with junction and intronic read quantitation in the *SpliceWiz* pipeline, resulting in minimal additional runtime and resource overhead. These optimizations allow *SpliceWiz* to seamlessly store and recall the data required to produce coverage plots on-demand.

### Coverage normalization of group replicates for visualization

Sequence coverage across alternatively spliced genomic regions differ between replicates of a biological group due to differences in sequencing depth and differential gene expression. Coverages of samples within a group can be combined by taking the group average of normalized individual per-nucleotide sequencing coverage. *SpliceWiz* uses transcript abundance (the sum of IR and spliced isoform abundances, see Supplementary Methods) to normalize coverage. This approach ensures that constitutively expressed elements (such as exons flanking the region containing the ASE) are normalized to unity.

## RESULTS

### SpliceWiz event filters improve PSI accuracy

To evaluate the effect of *SpliceWiz’s* event filters on accuracy of splicing quantitation, we simulated a dataset containing differentially expressed ASEs whereby ground truth PSI values can be determined from known transcript abundance values (Supplementary Methods). These transcript abundances were used to simulate sequencing reads in triplicates of two conditions using *flux-simulator version 1.2.1* [26]. We benchmarked *SpliceWiz*, with or without ASE event filters, alongside two other tools *rMATS turbo version 4.1.1* [11] and *SUPPA2 version 2.3* [27]. These tools were chosen as they are widely used and well-maintained tools that perform event-centric splicing quantitation. *rMATS* measures PSI using junction reads and reads that align exclusively to either included or excluded isoforms [11]. In contrast, *SUPPA2* measures PSI based on transcript expressions estimates provided by upstream tools [27].

We first determined optimal parameters for *SpliceWiz*’s event filters by assessing the performance of individual filters. For all *SpliceWiz* benchmarks, we uniformly applied a depth filter (Table S2A), given it is common practice for most tools to exclude low-expressing genes or transcripts. Additionally, we applied a participation or consistency filter at various levels of stringency using participation and consistency ratio cutoffs, respectively (Table S2A). We also assessed the effect of the terminal exon filter on alternate terminal event PSI accuracies. PSI accuracy performance was quantified as the area under the curve of the cumulative distribution function (AUC-CDF) of mean absolute errors of PSI for all measured ASEs. The benchmarks showed that the participation filter exhibited a mixed performance depending on the ASE modality (Figure 3A). The participation filter improved the accuracy most for SE events, with some improvement for A5SS/A3SS events, but worse accuracy for MXE and AFE/ALE events. The IR-participation filter led to slightly worse accuracy for IR events. In contrast, the consistency filter improved accuracy of both MXE and SE events (Figure 3B), with improvement of SE accuracy surpassing that of the participation filter. The IR-consistency filter improved the accuracy of IR events (Figure 3B). The terminal exon filter improved the accuracy of AFE/ALE events (Figure 3C).

**Figure 3:**
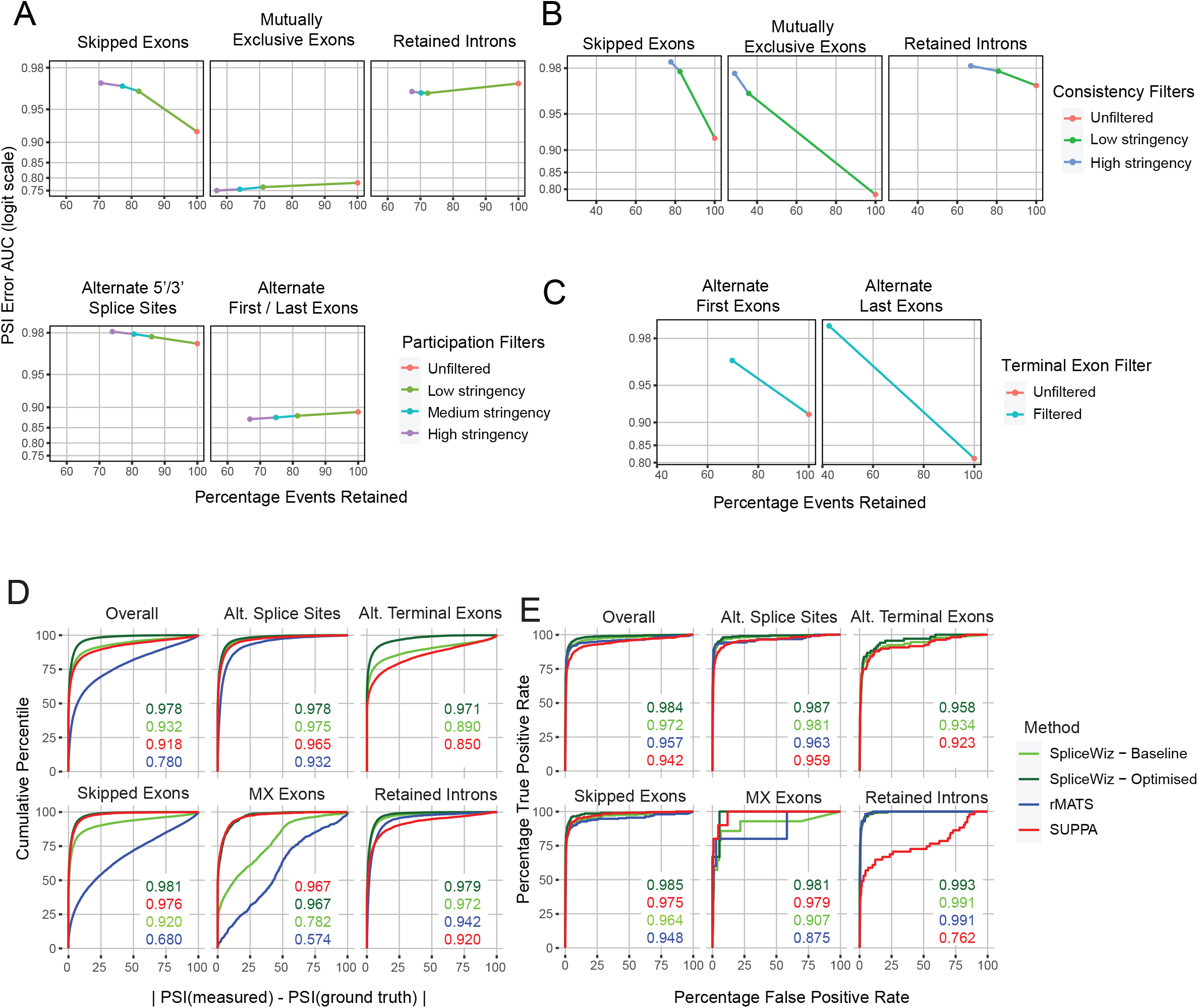
Percent-spliced-in measurement accuracy benchmarks of *SpliceWiz* against other tools. Accuracy benchmarks were performed using individual *SpliceWiz* filters **(A)** participation, **(B)** consistency and **(C)** terminal exon filters using the simulation dataset. The vertical axis is the area under the curve (AUC) of the cumulative distribution function of percent-spliced-in (PSI) mean absolute errors (PSI error AUC), as a surrogate measure of accuracy. The horizontal axis represents the percentage of alternative splicing events (ASE) after filtering, compared with the unfiltered set of ASEs. **(D)** Cumulative distribution function curves of PSI mean absolute errors of each tool in the PSI accuracy benchmark. Numbers indicate PSI error AUCs for each tool as displayed in their respective colors for each modality. **(E)** Modality-specific receiver operating characteristic (ROC) curves of differential ASE analysis. Area under the ROC curves are displayed for each tool as numbers in their respective colors for each ASE modality. Alt. Splice Sites = alternate 5’ / 3’ splice sites; Alt. Terminal Exons = alternate first / last exons; MX exons = mutually exclusive exons.

The best incremental gains in accuracy were observed at the lowest level of stringency of participation and consistency filters. Based on this, we defined an *optimized* set of filters consisting of low-stringency participation and consistency filters in combination with the terminal exon filter (Table S2B). Participation filters were not applied on MXE, AFE/ALE or IR events since they lowered accuracy. We benchmarked *SpliceWiz* with or without these optimized filters, alongside *rMATS* and *SUPPA2* each at default settings. For each tool, we included ASEs for evaluation if and only if finite PSI and p-values were returned by the evaluated tool.

Overall, we found that *SpliceWiz* using the baseline (depth) filter was more accurate at estimating PSI values than both *rMATS* and *SUPPA2* (Figure 3D). Further accuracy gains were seen when *SpliceWiz’s* optimized filters were applied (Figure 3D). PSI accuracy varied by the modality of ASE. *SUPPA2* was more accurate at measuring SE and MXE events compared with *rMATS* and baseline SpliceWiz. However, *SpliceWiz* with optimized filters surpassed *SUPPA2* at measuring SE events and reached a similar level of accuracy for MXE events. *SpliceWiz* performed the best for IR and A5SS/A3SS events. Both baseline and filtered *SpliceWiz* were more accurate than *SUPPA2* at measuring AFE/ALE events.

Next, we assessed the ability of each tool in detecting differential ASEs. Bias can arise for tools that use a different approach to annotate ASEs compared to that used to generate the simulation. Thus, for each tool, we constructed receiver-operator characteristic curves based only on their splicing annotations (Supplementary methods). We found that all 3 tools performed well at detecting differential ASE events (Figure 3E), although optimized *SpliceWiz* performed best overall. *rMATS* performed well at detecting differential SE events, despite its lower accuracy at estimating PSI. *SUPPA2* performed best in detecting differential MXEs but performed poorly at detecting differential IR compared to other tools.

We assessed the number of ASEs evaluated by each tool in the PSI accuracy benchmark. *SpliceWiz* evaluated more than 3 times the number of annotated IR events compared with *rMATS* and *SUPPA2* (Figure S4A), whereas for other forms of ASEs, *SpliceWiz* evaluated a similar number compared with *SUPPA2*. Optimized *SpliceWiz* filters resulted in a reduction of evaluated ASEs compared with baseline *SpliceWiz*. We then asked whether the differences in the number of evaluated ASEs arose from the different splicing annotations each tool generated from the input transcript annotation. When we tallied the number of ASEs in each tool’s splicing annotation, again we observed a 3-fold increase in IR events annotated by *SpliceWiz* compared with *rMATS* and *SUPPA2* (Figure S4B, C).

Because *rMATS* and *SUPPA2* produced a near-identical set of IR annotations, we scrutinised the methodology used by these two tools in defining IR events. In *SpliceWiz*, annotated IR events are defined simply as an intron in one transcript that lies within an exon of another transcript (Figure S3D). In contrast, *rMATS* and *SUPPA2* imposed a further criterion that two consecutive exons of one transcript (the exons flanking the putative retained intron) must have the same external boundaries as a single exon in another transcript (Figure S4D). Of the 15,188 extra retained introns annotated by *SpliceWiz*, 15,144 (99.7%) of these would have been excluded due to this extra criterion.

Taken together, *SpliceWiz* is an accurate and sensitive tool at determining and quantifying differential ASEs including IR. Event filters implemented in *SpliceWiz* resulted in increased measurement accuracy, especially for SE and MXE which feature isoforms with tandem junctions, and for AFE / ALE events. The improved IR annotation in *SpliceWiz* expands the number of assessed annotated IR events, thereby dramatically enhancing its sensitivity at detecting IR events.

### Parallel computation improves *SpliceWiz* performance

A major bottleneck to splicing quantitation is that most bioinformatics tools use a single CPU thread to process alignment data in BAM files. Additionally, some bioinformatic tools are implemented in higher-level programming languages (such as *R* and *python*) that greatly impair their performance [28]. To remedy these issues, we developed the *ompBAM* C++ API, which automates *OpenMP* based multithreaded BAM processing in *SpliceWiz* (Supplementary methods).

We tested the BAM processing performance of *SpliceWiz* alongside the three most popular *Bioconductor-based* tools for AS, namely *SGSeq version 1.24.0 [29], ASpli version 2.0.0* [30] and *IntEREst version 1.14.0* [16]. We also benchmarked the command line tools *rMATS version 4.1.1 (turbo) [11], MAJIQ version 2.2* [31] and *IRFinder version 1.3.1* [15]. Benchmarks were performed using BAM files from a downsampled version of the simulated dataset, comprising 6 samples each with approximately 25 million paired end reads. Four processor threads were used for all tools except *ASpli* (Supplementary methods).

We observed that *SpliceWiz* was the second fastest tool tested, with only *MAJIQ* being slightly faster (Figure 4A). In comparison with other tools available on *Bioconductor, SpliceWiz* was approximately 10 times faster than *ASpli*, 20 times faster than *SGSeq*, and 50 times faster than *IntEREst*. We conclude that the implementation of *SpliceWiz* in C++ greatly improves its computational performance, compared with *Bioconductor-based* tools. This performance is additionally enhanced by the multi-threaded implementation provided by *ompBAM*. As such, *SpliceWiz* is faster than most state-of-the-art command line tools.

**Figure 4:**
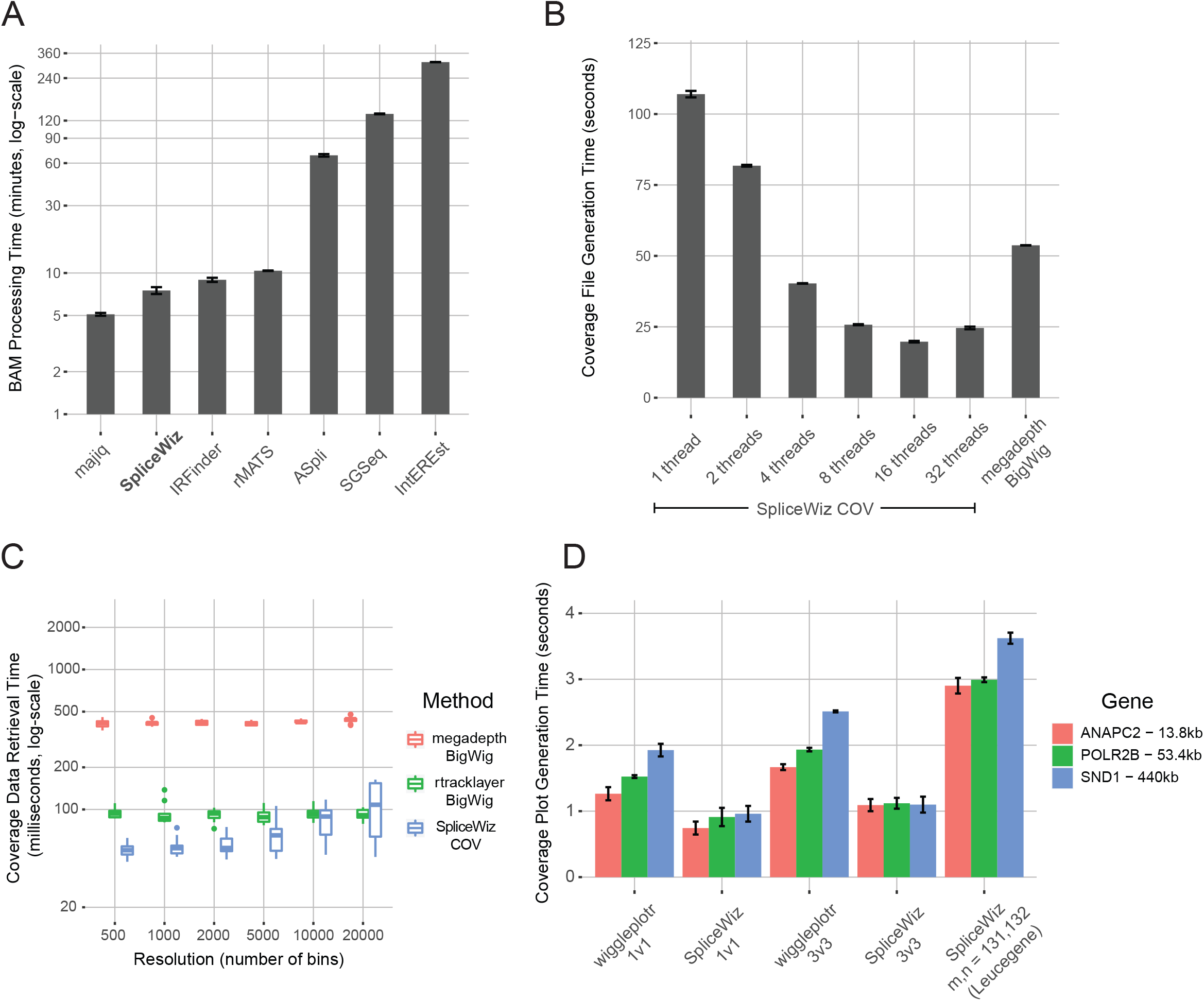
Performance benchmarks of *SpliceWiz*. **(A)** Mean BAM processing time of a 3-vs-3 dataset of BAM files (25 million paired-end reads per sample). For each tool benchmarked, 4 threads were used where multi-threading was supported. Error bars indicate standard deviation of triplicate runs. **(B)** Mean run time required to generate a single *COV* file (using *SpliceWiz*, with either single or multiple threads), compared with *BigWig* file generation using *megadepth*. Error bars indicate standard deviation of triplicate runs. **(C)** Mean retrieval time for RNA-seq coverage data of 10 randomly selected genes from *COV* and *BigWig* files. Resolution refers to the number of equal-sized bins the genomic region was divided for which the the mean coverage across each genomic bin was calculated. **(D)** Coverage plot generation mean runtime of *SpliceWiz* compared with *wiggleplotr*. For each tool, both 1-vs-1 (2 individual sample traces), and 3-vs-3 (triplicates of two conditions) were benchmarked. Additionally, *SpliceWiz* coverage plots of the Leucegene dataset split into two groups (n = 131, m = 132) were benchmarked. Error bars indicate standard deviation of triplicate runs.

### COV file format exhibits performance gains over BigWig-based tools

We designed the *COV* format as an alternative to the commonly used *BigWig* format to store strandspecific sequencing coverage, as well as to optimize data compression and computational performance in *SpliceWiz* (see Methods). Data compression performance was benchmarked by creating *BigWig* and *COV* files using *megadepth version 1.0.3* [32] and *SpliceWiz*, respectively. Data retrieval was benchmarked against the *Bioconductor* tools *megadepth* [32] and *rtracklayer version 1.50.0* [33], using *BigWig* and *COV* files constructed from the downsampled BAM files (25 million paired-end reads) of the simulation dataset. Plot generation time was benchmarked against the *Bioconductor* tool *wiggleplotr version 1.14.0* [34], which generates coverage plots from *BigWig* files.

First, we compared the sizes of *COV* and *BigWig* files derived from the BAM files. Relative to the parent BAMs, the mean COV file size was 1.56%, whereas that of *BigWig* was 1.97%. Considering that *COV* files store positive, negative as well as unstranded RNA-seq coverage data whereas *BigWig* stores only unstranded coverage data, *COV* files represent an estimated 2.5x gain in data compression compared with *BigWig* files.

Next, we examined the time taken to generate COV files. On a single thread, *COV* files take approximately twice as long to generate than *BigWig* files generated via the current fastest *BigWig* tool *megadepth* [32] (Figure 4B). However, *SpliceWiz COV* files can be generated using multiple threads. Using 8 threads, *COV* files are generated twice as fast compared to *BigWig* files (Figure 4B). Considering also that COV file generation in *SpliceWiz* is concurrent with splice junction and IR quantitation, *SpliceWiz’s COV* is computationally superior at generating RNA-seq coverage data.

We then benchmarked the computational performance in generating coverage plots. First, we assessed data retrieval in isolation. We measured the average data retrieval time of each of 10 randomly selected expressed genes in the simulation dataset. Coverage plots cannot represent data points of every nucleotide in their range as they are limited by pixel resolution (which is typically magnitudes lower than the number of nucleotides they represent). Due to this, horizontal and vertical positions of each pixel of the plot represent discrete genomic regions and their corresponding average coverage depth across these region, respectively. Thus, to perform resolution-specific benchmarks, the genomic region of each gene was divided into bins of equal size, the number of bins being the target horizontal resolution of the plot (between 500 and 10,000). Data retrieval for each tool involved computing the average coverage across each bin. This benchmark showed that, for resolutions up to 10,000 bins, *SpliceWiz* outperformed both *rtracklayer* and *megadepth*. It was on average 65% faster than *rtracklayer* and up to 7 times faster than *megadepth* (Figure 4C).

Finally, we measured the aggregate time required for plot generation, which includes both data retrieval and figure generation. *COV* and *BigWig* files generated from the simulation dataset were used to generate plots of three genes of different lengths (*ANAPC2*: 13.8 kb, *POLR2B*: 53.4 kb, and *SND1:* 440 kb). Individual (one versus one) and group-averaged (three versus three) coverage plots in *SpliceWiz* were benchmarked alongside *wiggleplotr*. In both comparisons *SpliceWiz* was approximately twice as fast (Figure 4D). Moreover, *SpliceWiz’s* performance did not decrease with increasing genomic window size, whereas *wiggleplotr* took longer to generate plots of longer genes. We also tested *SpliceWiz’s* coverage plots of the same genes on the much larger Leucegene dataset [35] (Gene Expression Omnibus accession GSE67039), which we split into two groups of 131 and 132 samples. Coverage plot generation took on average 3-4 seconds per plot (Figure 4D). Coverage plot generation via *wiggleplotr* was not feasible.

Taken together, *SpliceWiz’s* COV plot generation is superior to state-of-the-art *BigWig* implementations in both its computational speed and compression performance.

### Case study: Splicing factor expression correlates with differential alternative splicing in AML

*SpliceWiz* was designed to handle large experimental datasets of up to 1000 samples. To demonstrate its capabilities, we explored the correlation between splicing factor expression and differential ASEs in acute myeloid leukemia (AML). We processed aligned reads from 263 RNA-seq samples from the Leucegene AML dataset using *SpliceWiz*. We observed that, downstream to read alignment, the processing of BAM files is the longest step of the analysis, taking approximately 2.5 minutes per sample using 8 threads (Figure S5A).

After compilation and data import of isoform-specific counts, we evaluated each modality of ASEs differentially expressed in association with differential splicing factor expression. Using *SRSF10* as an illustrative example, we separated the dataset into 3 equal groups based on normalized gene counts of *SRSF10* (Figure S5B). After application of optimized ASE filters, differential ASE analysis was performed between the high and low *SRSF10* expressing groups. This was performed using both lognormal (*limma version 3.46.0* [19], Figure S5C) and beta-binomial (*DoubleExpSeq version 1.1* [20], Figure S6) distributions to model isoform counts. For each ASE modality, we counted differential ASEs as those that were statistically significant using both methods. The same analysis was repeated for the other 481 genes belonging to the “RNA splicing” gene ontology term (GO:0008380). Hierarchical clustering based on modality-specific differential ASEs were used to define 12 clusters of splicing factors (Figure 5A). These clusters are characteristic of their relative number of modality-specific ASEs associated with their differential expression (Figure 5B).

**Figure 5:**
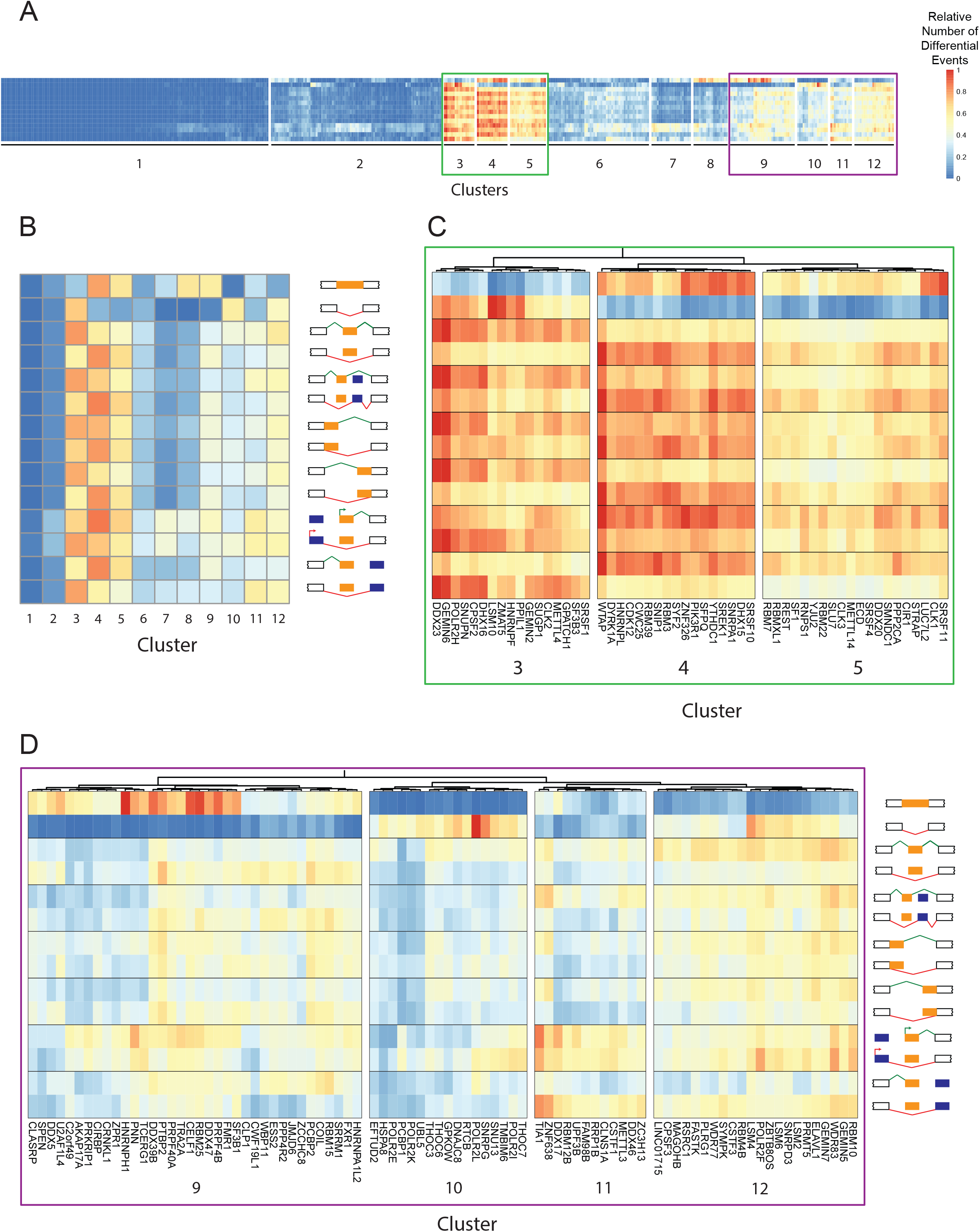
Alternative splicing signatures associated with splicing factor expression in AML. Differential alternative splicing event (ASE) analysis was performed between high and low expressing samples for each of 482 individual splicing factor genes. **(A)** Heatmap of differential ASEs (rows) between top and bottom third of samples (by splicing factor expression), for each of 482 splicing factors (columns). Heirarachical clustering identified 12 clusters of splicing factors. Quantities indicate the relative number of modality-specific differential ASEs associated with higher gene expression for each splicing factor. **(B)** Mean relative numbers of modality-specific differential ASEs for each cluster of splicing factors. **(C, D)** Enlarged heatmaps of clusters that exhibit high and moderate differential alternative splicing changes, respectively. See Figure S1 for descriptive labels of ASE modalities.

Of these, two groups of clusters are associated with high and moderate numbers of differential ASE (Figure 5C and 5D, respectively). Expression of splicing factors in cluster 3 was anti-correlated with IR, whereas those of clusters 4 and 5 exhibited the opposite pattern (Figure 5C). Interestingly, in clusters 3 and 4, IR was associated with exon skipping and usage of the downstream mutually exclusive exons. Additionally, IR was associated with splicing of longer introns in those with alternate splice sites. In contrast, the opposite pattern was observed in alternate first / last exon usage, whereby IR was associated with alternate terminal exon usage resulting in shorter introns. Less distinct patterns between ASE modalities were observed in splicing factor clusters 9-12 (Figure 5D).

Although it is difficult to identify the causative factors based on this correlative analysis, we conclude that differential expression of modality-specific ASEs provides useful observations that facilitate hypothesis generation regarding patterns of alternative splicing.

To demonstrate *SpliceWiz’s* ability to illustrate differential ASEs using coverage plots, we identified the top differentially expressed ASEs associated with higher *SRSF10* splicing factor expression. AML samples with high *SRSF10* expression have increased IR in *EXOSC10*, reduced inclusion of a skipped exon in *CTR9*, and increased inclusion of a mutually exclusive exon in *FYN* (Figure 6). We found that group-averaged coverage traces on the same plot provide an improved visualization of differential ASEs than traces of randomly chosen individual samples or group-averaged traces on separate plots (Figure S7A-C).

**Figure 6:**
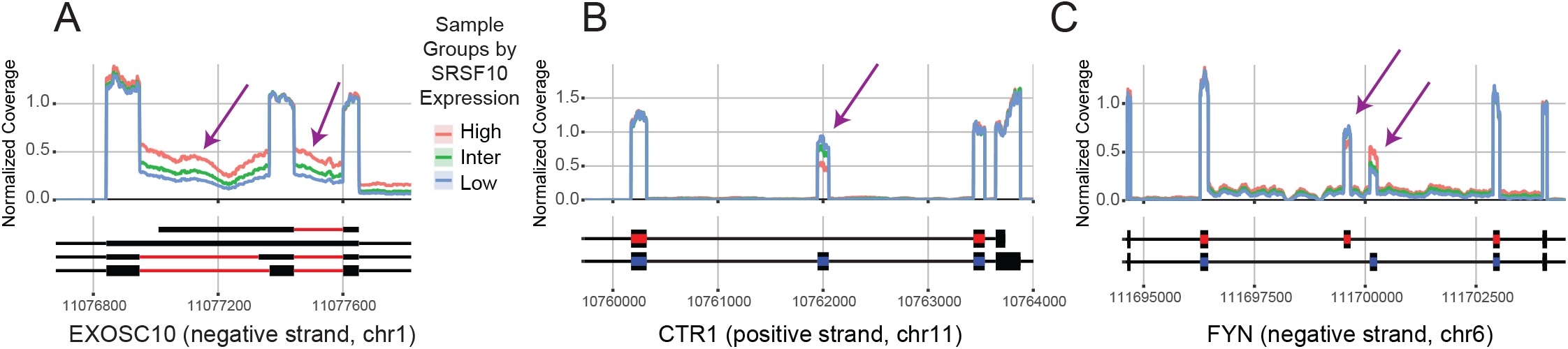
Example group-averaged coverage plots generated using *SpliceWiz*. Group-normalized coverage plots were generated using *SpliceWiz*, stratified by high, intermediate (inter) or low *SRSF10* expression. AML samples with high *SRSF10* expression have **(A)** increased intron retention in two tandem introns in *EXOSC10*, **(B)** reduced inclusion of the alternate exon in *CTR9*, and **(C)** increased inclusion of the upstream mutually exclusive exon in *FYN*. Arrows indicate regions of differential coverage due to group-wise differential splicing. On the genomic axis, blue and red rectangles indicate exons involved in the included and excluded isoforms, respectively, whereas red lines indicate retained introns.

Taken together, *SpliceWiz* provides a streamlined pipeline for quantifying and analyzing differential splicing from large datasets with potentially complex annotations. Group-normalized coverage plots provide effective, unbiased illustration of differential ASEs between experimental conditions.

## DISCUSSION

Here we demonstrated *SpliceWiz* as an ultra-fast and accurate graphical application that performs event centric AS quantitation, differential expression analysis and visualization. Differential ASEs between conditions can be visualized effectively using coverage plots that are appropriately normalized and averaged by experimental groups. *SpliceWiz* is an ideal application for evaluating alternative splicing at scale due to its computational efficiency and its ability to handle large numbers of samples. Its user-friendly graphical interface and its traditional and familiar annotation of alternative splicing as binary choices [9] enhances its accessibility to the general scientific audience. Additionally, we present *ompBAM* as an accessible and powerful API that automates multi-threaded processing of alignment BAM files for other R package developers.

Transcriptomic complexity, as occurs in higher eukaryotes including human and mouse, results in numerous regions of genomic overlap between transcript isoforms. Accordingly, attribution of short sequence reads to individual transcript isoforms is often impossible. Event-based splicing quantitation is an attractive approach, by assigning junction reads to transcript groups based on inclusion and exclusion of alternative exon regions (and introns in the case of IR). However, as we have shown, SE and MXE events are often inaccurately measured because they involve two consecutive junction reads. Their measurement is often confounded by expression of isoforms including one but not the other junction. *MAJIQ* addresses this by forgoing tandem junction counts; instead, it quantifies *local splicing variations* [31]. The complexity of overlapping ASEs is addressed formally in *Whippet* as *entropy*. However, its authors showed that higher *entropy* events are less accurately quantified [36]. In *SpliceWiz*, we addressed transcriptomic complexity using ASE filters to identify the most readily measurable ASEs based on the data. We showed that this approach increases measurement accuracy at the expense of a modest decrease in measurable ASEs.

However, for some ASE modalities, we observed divergent performances between PSI measurement accuracy and detection accuracy of differential ASEs. For example, *rMATS* performed poorly at estimating PSIs for SE events whereas it performed well at detecting differential SE events. This indicates that the inaccurate PSI estimates were systematic errors that canceled out when calculating differential PSIs. Another unexpected observation is that the IR-participation filter led to lower PSI accuracy in the simulation dataset. The IR-participation filter removes IR events with a low proportion of the intron covered by reads, effectively removing events in which IR is not occurring. In effect, this filter removes events that can be accurately measured (as PSI equals zero in all conditions) but which are not differentially expressed. We posit that, although the IR-participation results in lower PSI measurement accuracy, it may improve differential IR detection accuracy in real-world scenarios by removing events in which there is a high degree of confidence that IR is absent.

A limitation of *SpliceWiz* is that it only assesses annotated ASEs, except in IR where both novel and annotated IR events are assessed. Given the average human exon is approximately 120 nucleotides long [37], the exact combination of exons of a transcript derived from unannotated ASEs cannot be readily determined from short read sequencing. However, novel ASEs are of biological interest as they frequently feature in malignancies [38, 39]. One approach to quantify novel ASEs using *SpliceWiz* would be to use long-read sequencing (of representative samples) to generate or supplement *SpliceWiz’s* annotation of ASEs. This approach would have the added advantage of reducing transcriptomic complexity by removing the annotation of transcripts that are not represented in the dataset.

Genome alignment, considered more accurate albeit more computationally demanding than pseudo-alignment [40], requires storage of alignment data in compressed format as BAM files. Accessing this data is not a trivial programmatic task, and as such, most R package developers rely on low-level APIs such as the C-based *htslib* library [14] and the high-level *RSamtools* R package [13]. Performance of tools that use *htslib* and *RSamtools* can be bottlenecked due to interfacing with high-level languages such as R and Python, and the lack of multi-threading support. Thus, developers wishing to utilize multiple threads to access BAM files must either multi-thread file access (one file per thread), or implement their own multi-threaded BAM processing from scratch. The *ompBAM* package, upon which *SpliceWiz* is built, offers developers a well-documented API to write C++ based functions using the *Rcpp* environment [21]. These functions can be exported to the R environment to be used by the encompassing R package. In addition to gene expression and splicing analysis, *ompBAM* can accelerate processing of BAM files used for DNA methylation analysis, SNP analysis, as well as long-read sequencing. We anticipate that the development of *ompBAM*-based R packages will lead to an expansion of computationally efficient bioinformatic tools.

RNA sequencing coverage plots, in conjunction with sashimi plots [12], are a popular way to visualize ASEs. However, most implementations are limited by their inability to compare across experimental groups, and inflexibility with regards to strand-specific coverage. Here, we show that group differences in percent-spliced-in ratios can be clearly visualized using coverage plots normalized and aggregated by experimental groups. *SpliceWiz’s* implementation of interactive coverage plots is feasible because of its computational efficacy, due in part to its use of a new data format to store strand-specific coverage. Group-wise coverage plots are not entirely novel. *Manananggal* previously implemented a similar visualization for exon-centric differential expression [41]. However, *SpliceWiz* also visualizes intronic coverage thereby allowing for visualization of IR. Group-wise averaging in SpliceWiz is performed using a normalization factor which ensures constitutively-spliced flanking elements are normalized to unity. One limitation to coverage-based ASE visualization is that differential ASEs with high inclusion rates (i.e., PSI approaching 1) are poorly visualized. This is likely because the higher inter-sample variations in coverage, which accompany high inclusion ratios, conceal the comparatively low coverage difference in such ASEs.

Finally, *SpliceWiz* provides a GUI which enables user-friendly point-and-click functionality. Although other tools such as *psichomics* [42] provide GUIs, *SpliceWiz* enables interactive exploration by allowing users to select ASEs of interest through tabular data, volcano and scatter plots. Selected ASEs can be visualized using heatmaps of PSIs and coverage plots. Such interactive data exploration methods are under-utilized in current bioinformatic tools and can vastly improve the user experience.

In conclusion, we provide *SpliceWiz* as a software suite for differential splicing analysis. SpliceWiz harnesses efficient multi-threaded processing of alignment data through its use of the *ompBAM* API. Innovations include novel filtering approaches to enhance ASE measurement and detection accuracy; a file format to efficiently store and recall stranded coverage data; and group-averaged coverage plots to compare differential splicing between experimental conditions. Its graphical interface and interactive visualization vastly improves the user experience and its performance optimizations streamline the analyses of large datasets. *SpliceWiz* will accelerate discovery research in the field of alternative splicing.

## DATA AVAILABILITY

*SpliceWiz* and *ompBAM* are available on Bioconductor. The developmental version of *SpliceWiz* is available on GitHub (https://github.com/alexchwong/SpliceWiz). Flux simulator input files and *SpliceWiz* output files of the simulation dataset, can be found at https://github.com/alexchwong/SpliceWizResources.

## SUPPLEMENTARY DATA

Supplementary Data are available as a separate file.

## ACKNOWLEDGEMENT

The authors acknowledge the Sydney Informatics Hub and the use of the University of Sydney’s high performance computing cluster, Artemis.

## FUNDING

This work was supported by the National Health and Medical Research Council (Grant #2010647 to AW and JJLW, #1128175 to JJLW and JEJR, #1177305 to JEJR and #1196405 to US); Cure the Future (JEJR); the Cancer Council NSW (Grant # RG20-12 to US); the Tropical Australian Academic Health Centre (Grant # SF0000321 to US) and an anonymous foundation (JEJR). ACHW was a recipient of the Australian Government Research Training Program PhD Scholarship.

## CONFLICT OF INTEREST

The authors declare that they have no conflict of interest.

## Notes

### Competing Interest Statement

The authors have declared no competing interest.

## REFERENCES

1. Wahl MC, Will CL, Lührmann R. The spliceosome: design principles of a dynamic RNP machine. Cell. 2009;136(4):701–18.

2. Reed R. Initial splice-site recognition and pairing during pre-mRNA splicing. Curr Opin Genet Dev. 1996;6(2):215–20.

3. Fiszbein A, Kornblihtt AR. Alternative splicing switches: Important players in cell differentiation. Bioessays. 2017;39(6).

4. Lin JC, Tsao MF, Lin YJ. Differential impacts of alternative splicing networks on apoptosis. Int J Mol Sci. 2016;17(12):2097.

5. Shalgi R, Hurt JA, Lindquist S, Burge CB. Widespread inhibition of posttranscriptional splicing shapes the cellular transcriptome following heat shock. Cell Rep. 2014;7(5):1362–70.

6. Schmid R, Grellscheid SN, Ehrmann I, Dalgliesh C, Danilenko M, Paronetto MP, et al. The splicing landscape is globally reprogrammed during male meiosis. Nucleic Acids Res. 2013;41(22):10170–84.

7. Wong JJ, Ritchie W, Ebner OA, Selbach M, Wong JW, Huang Y, et al. Orchestrated intron retention regulates normal granulocyte differentiation. Cell. 2013;154(3):583–95.

8. Boutz PL, Bhutkar A, Sharp PA. Detained introns are a novel, widespread class of post-transcriptionally spliced introns. Genes Dev. 2015;29(1):63–80.

9. Blencowe BJ. Alternative splicing: new insights from global analyses. Cell. 2006;126(1):37–47.

10. Vanichkina DP, Schmitz U, Wong JJ, Rasko JEJ. Challenges in defining the role of intron retention in normal biology and disease. Semin Cell Dev Biol. 2018;75:40–9.

11. Shen S, Park JW, Lu Z-x, Lin L, Henry MD, Wu YN, et al. rMATS: Robust and flexible detection of differential alternative splicing from replicate RNA-Seq data. Proceedings of the National Academy of Sciences. 2014;111(51):E5593.

12. Katz Y, Wang ET, Airoldi EM, Burge CB. Analysis and design of RNA sequencing experiments for identifying isoform regulation. Nat Methods. 2010;7(12):1009–15.

13. Danecek P, Bonfield JK, Liddle J, Marshall J, Ohan V, Pollard MO, et al. Twelve years of SAMtools and BCFtools. Gigascience. 2021;10(2).

14. Bonfield JK, Marshall J, Danecek P, Li H, Ohan V, Whitwham A, et al. HTSlib: C library for reading/writing high-throughput sequencing data. Gigascience. 2021;10(2).

15. Middleton R, Gao D, Thomas A, Singh B, Au A, Wong JJ, et al. IRFinder: assessing the impact of intron retention on mammalian gene expression. Genome Biol. 2017;18(1):51.

16. Oghabian A, Greco D, Frilander MJ. IntEREst: intron-exon retention estimator. BMC Bioinformatics. 2018;19(1):130.

17. Li HD, Funk CC, Price ND. iREAD: a tool for intron retention detection from RNA-seq data. BMC Genomics. 2020;21(1):128.

18. Love MI, Huber W, Anders S. Moderated estimation of fold change and dispersion for RNA-seq data with DESeq2. Genome Biol. 2014;15(12):550.

19. Ritchie ME, Phipson B, Wu D, Hu Y, Law CW, Shi W, et al. limma powers differential expression analyses for RNA-sequencing and microarray studies. Nucleic Acids Res. 2015;43(7):e47.

20. Ruddy S, Johnson M, Purdom E. Shrinkage of dispersion parameters in the binomial family, with application to differential exon skipping. The Annals of Applied Statistics. 2016;10(2):690–725.

21. Eddelbuettel D, Francois R. Rcpp: Seamless R and C++ integration. Journal of Statistical Software. 2011;40(8):1 – 18.

22. Morgan M, Shepherd L. AnnotationHub: Client to access AnnotationHub resources. R package version 3.2.1. 2022.

23. Dobin A, Davis CA, Schlesinger F, Drenkow J, Zaleski C, Jha S, et al. STAR: ultrafast universal RNA-seq aligner. Bioinformatics. 2013;29(1):15–21.

24. Liao Y, Smyth GK, Shi W. The R package Rsubread is easier, faster, cheaper and better for alignment and quantification of RNA sequencing reads. Nucleic Acids Research. 2019;47(8):e47–e.

25. Kim D, Paggi JM, Park C, Bennett C, Salzberg SL. Graph-based genome alignment and genotyping with HISAT2 and HISAT-genotype. Nature Biotechnology. 2019;37(8):907–15.

26. Griebel T, Zacher B, Ribeca P, Raineri E, Lacroix V, Guigó R, et al. Modelling and simulating generic RNA-Seq experiments with the flux simulator. Nucleic Acids Res. 2012;40(20):10073–83.

27. Trincado JL, Entizne JC, Hysenaj G, Singh B, Skalic M, Elliott DJ, et al. SUPPA2: fast, accurate, and uncertainty-aware differential splicing analysis across multiple conditions. Genome Biology. 2018;19(1):40.

28. Aruoba SB, Fernández-Villaverde J. A comparison of programming languages in economics. National Bureau of Economic Research Working Paper Series. 2014;No. 20263.

29. Goldstein LD, Cao Y, Pau G, Lawrence M, Wu TD, Seshagiri S, et al. Prediction and quantification of splice events from RNA-Seq data. PLoS One. 2016;11(5):e0156132.

30. Mancini E, Rabinovich A, Iserte J, Yanovsky M, Chernomoretz A. ASpli: integrative analysis of splicing landscapes through RNA-Seq assays. Bioinformatics. 2021;37(17):2609–16.

31. Vaquero-Garcia J, Barrera A, Gazzara MR, González-Vallinas J, Lahens NF, Hogenesch JB, et al. A new view of transcriptome complexity and regulation through the lens of local splicing variations. Elife. 2016;5:e11752.

32. Wilks C, Ahmed O, Baker DN, Zhang D, Collado-Torres L, Langmead B. Megadepth: efficient coverage quantification for BigWigs and BAMs. Bioinformatics. 2021;37(18):3014–6.

33. Lawrence M, Gentleman R, Carey V. rtracklayer: an R package for interfacing with genome browsers. Bioinformatics. 2009;25(14):1841–2.

34. Alasoo K. wiggleplotr: Make read coverage plots from BigWig files. R package version 1.15.0. 2021.

35. Lavallée VP, Baccelli I, Krosl J, Wilhelm B, Barabé F, Gendron P, et al. The transcriptomic landscape and directed chemical interrogation of MLL-rearranged acute myeloid leukemias. Nat Genet. 2015;47(9):1030–7.

36. Sterne-Weiler T, Weatheritt RJ, Best AJ, Ha KCH, Blencowe BJ. Efficient and Accurate Quantitative Profiling of Alternative Splicing Patterns of Any Complexity on a Laptop. Mol Cell. 2018;72(1):187–200.e6.

37. Mokry M, Feitsma H, Nijman IJ, de Bruijn E, van der Zaag PJ, Guryev V, et al. Accurate SNP and mutation detection by targeted custom microarray-based genomic enrichment of short-fragment sequencing libraries. Nucleic Acids Res. 2010;38(10):e116.

38. Monteuuis G, Schmitz U, Petrova V, Kearney PS, Rasko JEJ. Holding on to junk bonds: intron retention in cancer and therapy. Cancer Research. 2021;81(4):779–89.

39. Wang BD, Lee NH. Aberrant RNA splicing in cancer and drug resistance. Cancers (Basel). 2018;10(11):458.

40. Wu DC, Yao J, Ho KS, Lambowitz AM, Wilke CO. Limitations of alignment-free tools in total RNA-seq quantification. BMC Genomics. 2018;19(1):510.

41. Barann M, Zimmer R, Birzele F. Manananggal - a novel viewer for alternative splicing events. BMC Bioinformatics. 2017;18(1):120.

42. Saraiva-Agostinho N, Barbosa-Morais NL. psichomics: graphical application for alternative splicing quantification and analysis. Nucleic Acids Research. 2019;47(2):e7.

