## Supplementary Methods, Tables and Figures for "SpliceWiz: easy, optimized, and accurate alternative splicing analysis in R"

#### Supplementary Data

#### Supplementary Methods

##### Implementation of multi-threaded file processing in ompBAM

*ompBAM* is an R package that includes a set of C++ header files for R package developers. It also contains R-based functions that facilitate new package development, a detailed vignette, documentation, as well as a fully functional example package.

Developers can use *ompBAM* in their C++ based functions via the *Rcpp* API [1] by linking to the main *ompBAM* header file. BAM file processing is handled by the *pbam_in* object, while processing of alignments is handled by the *pbam1_t* object.

Internally, the *pbam_in* object decompresses the BAM file using the *zlib* library provided by the *zlibbioc* R package. File input is performed either using one or multiple threads. When one thread is used for file input, the remaining threads are used for data decompression which occurs asynchronously. In *SpliceWiz*, one thread is used for file decompression by default.

File input is stored into memory using a primary and secondary buffer, each of equal size. The first time the body of the BAM file is accessed, compressed data is read from the file and stored in the primary buffer as well as in a pre-determined fraction (hereafter *chunk*) of the secondary buffer. When more than one *chunk's* worth of data from the primary buffer is decompressed, subsequent file read is triggered whereby another *chunk* of data is read from file and stored in the secondary buffer. When less than one *chunk's* worth of data in the primary buffer remains, a buffer swap occurs. This buffer swap consists of 4 steps: (A) transfer of the remaining data from the primary buffer to its first position; (B) transfer of most of the secondary buffer into the remainder of the primary buffer; (C) transfer of the residual data of the secondary buffer to its first position; and (D) file input to fill the secondary buffer up to one *chunk's* worth of data.

Decompression is performed using multiple threads and can occur thread-asynchronously with file access (when one-thread is used for file access) or after file access (when multiple threads are used for file access). Decompressed alignment data is stored in a single allocated memory buffer and is divided into contiguous blocks by the number of available threads.

When initializing the *pbam_in* object, users specify the desired number of CPU threads, the size of the file and decompressed data buffers, the *chunk* fraction (the ratio of the chunk size to that of the file buffer), and whether to access the file with one or with multiple threads. *SpliceWiz* uses the *ompBAM* default settings, which is 500 Mb for each of the two file buffers, 1 Gb decompressed data buffer, and a chunk fraction of 5. A call to the *fillReads()* function instructs *pbam_in* to perform file input and decompression as described above. After this, the *supplyRead()* function is called to retrieve alignments from the decompressed data. *supplyRead()* is thread-safe, and is designed to be called from within a parallel FOR loop using *OpenMP*. *supplyRead()* returns a *pbam1_t* object that acts as a virtual pointer to the memory storing the decompressed data. Handling alignments as virtual pointers helps improve performance by minimizing *memcpy* calls. *pbam1_t* contains functions that retrieve alignment data, including read name, cigar, sequence, and data contained within tags.

**Quantifying Percent-Spliced-In (PSI) and transcript abundance**

ASEs, including annotated IR events, are quantified in *SpliceWiz* using PSI, which is calculated using equation 1. $Incl$ and $Excl$ are the estimated abundances of included and excluded isoforms, respectively. For definitions of how *SpliceWiz* annotates included and excluded isoforms, see Figure S1.

$$(1)\ \ \mathrm{PSI} = \frac{\mathrm{Incl}}{\mathrm{Incl} + \mathrm{Excl}}$$

For IR, $Incl$ is calculated using the trimmed mean depth of coverage across the measured intron; otherwise, $Incl$ and $Excl$ are calculated by summing isoform-specific junction reads. For SE, $Incl$ is the average junction reads of the two junctions flanking the included intron; for MXE, both $Incl$ and $Excl$ are determined from the average junction reads flanking the mutually exclusive exons.

*SpliceWiz* separately determines IR-ratios of all introns (regardless of annotation). For annotated IR events, SpliceWiz calculates and performs differential analysis using both PSI and IR-ratio, whereas differential IR-ratio is only performed on novel IR events. IR-ratio is defined by equation 2, where $IRT$ and $Spl$ are estimated abundances of IR and spliced transcripts, respectively.

$$(2)\ \ \mathrm{IR_{ratio}} = \frac{\mathrm{IRT}}{\mathrm{Transcript\ Abundance}} = \frac{\mathrm{IRT}}{\mathrm{IRT} + \mathrm{Spl}}$$

In contrast to PSI (where $Excl$ refers only to junction reads specific to the intron of interest), $Spl$ refers to the sum of junction reads across the intron of interest, as well as those of overlapping introns. Thus, whereas PSI is measured with an implicit assumption that the expressed transcripts can only either contain the spliced or retained intron, IR-ratio accounts for the expression of transcripts arising from splicing of overlapping introns. IR-ratio differs most from PSI when the intron of interest belongs to the minor isoform in a sample, where $Excl \ll Spl$ , thereby PSI can result in overestimation of IR compared with IR-ratio. Therefore, the method used to calculate $Spl$ to appropriately account for overlapping spliced transcripts is crucial in accurately determining IR-ratio.

*IRFinder* [2] calculates $Spl$ by summing junction reads where at least one flanking exon shares a splice site with the intron of interest (in *IRFinder* this parameter is referred to as *SpliceMax)*. However, IR overestimation may still occur in cases where the major isoform involves flanking exons where neither exon shares splice sites with the intron of interest (Figure S8A, B). To address this in *SpliceWiz*, we defined $Excl$ using the *SpliceOver* parameter to encompass junction counts of such isoforms. *SpliceOver* is determined using junction counts not only across flanking exons, but also across flanking *exon clusters*. These *exon clusters* are regions of alternate exons that overlap the exons flanking the intron of interest (Figure S8A). We found that in some IR events, *SpliceWiz's SpliceOver* metric led to lower IR-ratios compared with *IRFinder's SpliceMax* (Figure S8B). This confirmed that *SpliceOver* corrects the overestimation of IR due to its inclusion of junction reads overlapping the intron of interest.

In addition to calculating IR-ratios, transcript abundance (the sum of $IRT$ and $Spl$) is also used as the normalization parameter for group-averaged coverage plots. In other words, for each sample, pernucleotide coverages are divided by transcript abundance. This ensures the exon abundances of flanking exons are normalized to unity in each sample, prior to group-averaging. Using transcript abundance for both IR-ratio calculation and coverage normalization ensures that the visualized intron coverage depths accurately represent the calculated IR-ratios.

##### Determining Participation and Consistency Ratios

Participation ratio is the summed expression of both isoforms as a proportion of all isoforms with junction reads bridging the same exon clusters, as shown in equation 3 (where $a_j$ and $a_k$ represent the expression of the two isoforms of the ASE of interest, and $a_1$, …, $a_n$ are the expression of $n$ possible isoforms sharing the exon clusters from which the ASE originates).

$$(3)\ \ Participation_{ratio} = \frac{a_j + a_k}{\sum_{i=1}^{n} a_i}$$

Consistency ratio is applied to isoforms that comprise tandem junctions. It refers to the ratio of the counts of reads aligned to the less numerous tandem junction, to that of the more numerous, as shown in equation 4 (where $j$ and $k$ are the counts of the two tandem junctions).

$$(4)\ \ Consistency_{ratio} = \frac{min\ (j,k)}{max\ (j,k)}$$

Participation and consistency ratios are used as thresholds to filter ASEs (see Supplementary Table S2A,B).

##### Annotation of alternative splicing events in *SpliceWiz*

In *SpliceWiz*, ASEs are annotated by the set of coordinates of splice junctions defining included or excluded isoforms (Figure S1). In principle, SpliceWiz defines the included isoform as the one that “includes” a larger proportion of the gene or that which contains the shorter upstream intron. Specifically, the included isoform is defined as:

- the included exon (in SE)
- the 5’-most mutually exclusive exon (in MXE)
- the retained intron (in IR)
- the shorter spliced intron (in A5SS, A3SS, AFE, ALE).

To identify SE and MXE events, for each transcript a list of skip junctions [$D_i$, $A_{i+1}$] is compiled. $D$ and $A$ denote donor and acceptor splice junction coordinates; $i$ representing the intron number in the transcript; $i \in \{1, \ldots, n-1\}$ where $n$ is the number of introns in the transcript. SE events were defined by matching a skip junction of the included isoform with an intron of the excluded isoform. MXEs were defined by unique transcripts sharing a common skip junction but containing cassette exons [$A_i$, $D_{i+1}$] with different 5’- and 3’-coordinates.

AFEs were defined as transcript pairs with first introns sharing a common acceptor site, whereas ALEs were defined as transcript pairs with last introns sharing a common donor site. A5SS and A3SS events were defined as transcript pairs sharing introns with common acceptors and donors, respectively. Additionally, alternative splice sites in A5SS and A3SS events must belong to the same exon cluster and must be distinct from AFE or ALE events.

*SpliceWiz* compiles a list of both annotated and unannotated IR events. All introns of transcripts belonging to transcript biotype categories *protein_coding*, *processed_transcript*, *lincRNA*, *antisense* and *nonsense_mediated_decay* were considered as eligible IR events. An annotated IR event was defined as an eligible IR event for which an exon from another transcript (belonging to any transcript biotype category excluding *sense_intronic* and *retained_intron*) spans its entire length (Figure S2D). An unannotated IR event was defined as all eligible IR events that were not defined as annotated IR events.

Optionally, users can choose to reduce the number of tested unannotated IR events by filtering overlapping introns. This is performed by determining the mean junction reads aligned to introns, and iteratively removing overlapping introns with lower expression. Removal of overlapping introns prevents their double counting.

##### Differential alternative splicing event analysis in SpliceWiz

In *SpliceWiz*, differential ASE analysis is performed by modelling counts of inclusion and exclusion isoforms using log-normal, negative binomial, or beta binomial distributions. The statistical modelling is performed using wrappers to the relevant functions from the R packages *limma* [3], *DESeq2* [4], and *DoubleExpSeq* [5], respectively. Inclusion and exclusion isoform counts for all ASE modalities, except IR, are counts of reads aligned across splice junctions. For inclusion counts of IR, we adopted the method used in *IRFinder* [2]. Specifically, it is the trimmed mean of the depth of intron coverage across the measurable intron. The measurable intron was defined as the genomic region occupied by the intron, excluding the first and last 5 nucleotides, regions occupied by partially overlapping exons of non-IR transcripts (plus 5 flanking nucleotides on both ends), and regions of low mappability. *SpliceWiz* uses the interaction term between inclusion and exclusion counts to model the effect of user-defined experimental conditions on differential ASE expression. For example, in *limma*, to model an experiment by a user-defined test variable *Treatment*, with two batch factors *Batch1* and *Batch2*, would be:

$$design = Batch1 + Batch2 + Treatment + Treatment{:}Isoform$$

where *Isoform* is a categorical variable to indicate whether the counts are of inclusion or exclusion isoforms. To assess differential ASE between two conditions *(A)* and *(B)* as specified by *Treatment*, the contrasting terms *Treatment(A):Isoform(Included)* and *Treatment(B):Isoform(Included)* would be used. An analogous method is used for *DESeq2*. Additionally, for *DESeq2*, the test variable can be treated as a continuous variable, allowing for time series analysis. In *DoubleExpSeq,* contrasts between two conditions (i.e., without normalization for batch factors) is implemented.

##### Simulated dataset

We sought to create a simulated dataset based on pre-determined transcript expression profiles. In order to create expression profiles resembling that of biological samples, we first quantified transcript expressions of the THP-1 (undifferentiated) cell line from Green *et al* [6] (GSE130011) using *SALMON version 1.4.0* [7], the human reference genome (GRCh38/hg38) and gene annotations from Ensembl release-94. We calculated the means and dispersions of these transcript expression profiles using *DESeq2 version 1.30.1* [4], and used these to seed initial transcript expression values in triplicates of both control and treatment groups based on the negative binomial distribution with moderated dispersion.

We first tested whether this random initialization of transcript expression values resulted in differential ASEs. We determined ground-truth differential alternative splicing events (ASEs) by assuming that logit-transformed percent-spliced-in (PSI) values were normally distributed. Statistical significance was determined using Student's t-test on logit-transformed PSIs. We observed that statistically significant false positive ASEs (as defined as having an adjusted p value < 0.05, using the Benjamini Hochberg method) never featured a minor isoform mean expression higher than 0.5 transcripts per million (tpm). We thus defined ASEs as true positive differential ASEs if it satisfied the following criteria:

- Absolute difference in mean PSIs of at least 0.05 between control and treatment groups,
- Adjusted p value of 0.05 or less,
- Mean expression of the minor isoform (across the dataset) of at least 0.5 tpm

To generate true positive differential ASEs, we randomly selected one annotated ASE belonging to each of 500 randomly selected genes. For each of these ASEs, included and excluded isoform expressions were increased or decreased in the treatment group, while keeping overall gene expression constant, such that its differential ASE expression would be statistically significant between treatment and control groups (adjusted p value < 0.05). We then re-analyzed the transcript expressions and identified 383 ground-truth differentially expressed ASEs (A5SS: 85, A3SS: 51, AFE: 40 ALE: 8, MXE: 7, RI: 59, and SE: 129).

Finally, using the simulated transcript expression profiles, we simulated 100 million paired-end reads of 100 nucleotides in length, per replicate, using flux-simulator (25). The parameters used in *flux-simulator* are indicated in Table S3.

**Experimental dataset**

263 samples from the Leucegene dataset were obtained in SRA format from the NCBI Gene Expression Omnibus [8] (GSE67039) and converted to FASTQ using *sratoolkit* version 2.10.0.

**Alignment and analysis of experimental and simulated datasets**

Raw paired-end sequences of experimental and simulated datasets were aligned to the human genome (GRCh38/hg38) using *STAR* aligner *version 2.7.3a* [9]. Gene counts were obtained using the *featureCounts* method in the *Rsubread* R package *version 2.4.3* [10], and normalized gene counts were generated using the *voom* function of the *limma* R package *version 3.46.0*. Differential ASE analysis was performed using *SpliceWiz* (using its *DoubleExpSeq* differential ASE function, *DoubleExpSeq version 1.1*) and *rMATS* using the alignment BAM files. For differential ASE analysis of the Leucegene dataset, both *DoubleExpSeq* and *limma* differential ASE functions were used. Transcript expression profiles obtained using *SALMON* (using the raw sequences as input) were used for *SUPPA2* analysis. Ensembl release-94 gene annotations were used in *Rsubread*, *SpliceWiz*, *rMATS* and *SALMON*.

**Benchmarks of accuracy**

Ground-truth PSI values were calculated based on the pre-determined transcript expression values in the simulated dataset. These were compared with measured PSI values from *SpliceWiz*, *rMATS* and *SUPPA2* to determine mean absolute error of PSI values. Accuracy of PSIs was determined by calculating the area under the curve of the cumulative distribution function of mean absolute error PSI values.

Accuracy of differential ASE analysis was determined by calculating the area under the receiver-operator characteristic curve (AUROC). ROC curves were generated using *ROCit version 2.1.1*. For each of *SpliceWiz*, *rMATS* and *SUPPA2*, nominal p values were used to rank differential ASEs. We sought to avoid biases arising from the different ASE annotations generated by each tool, as well as false positive differential ASEs arising from low expression of the minor ASE isoform. Thus, we excluded ASEs that are not included in the annotation of the tool being benchmarked, and all ASEs with ground-truth minor isoform expression less than 0.5 tpm.

**Benchmarks of computational performance**

Processing time of BAM files were benchmarked using the BAM files of the simulated dataset. Benchmarks were performed using 4 processor threads in all tools except *ASpli*, where there was no option to enable multi-threading. For *IRFinder*, one processor thread was used for *gzip*-based data decompression while a second thread was used to process alignments; thus 2 processor threads per sample were allowed. The parameters used to benchmark each tool is as indicated in Table S4.

COV file generation time and file size was benchmarked by generating COV files using *SpliceWiz*, using the *BAM2COV()* command. This was compared with that of *BigWig*, generated using *megadepth* using the command *bam_to_bigwig()*.

Data retrieval time was benchmarked between *COV* and *BigWig* formats. We determined expressed genes in the simulated dataset using *Rsubread's featureCounts*() function, excluding genes for which the mean read counts were less than 1000. We randomly selected 10 genes from this filtered list. For each of these 10 genes, we recorded the time taken to retrieve the coverage data from *COV* and *BigWig* files. COV file data retrieval was performed using *SpliceWiz's GetCoverageBins()* function. *BigWig* data retrieval was performed using *megadepth's get_coverage()* function. Additionally, we used the *import.bw()* function of the *rtracklayer* package, and additional custom code such that equivalent data was returned.

Plot generation time was benchmarked between *COV* and *BigWig* formats using *SpliceWiz* and *wiggleplotr*, respectively. For both, the *plotCoverage*() function (identically named in both packages) was benchmarked.

The *microbenchmark* R package *version 1.4-7* was used to measure all time-related benchmarks. All computation benchmarks (except processing of the Leucegene dataset) were performed on a Linux server. BAM and COV files were stored in page cache prior to benchmarking, using the *vmtouch* utility. Performance observation of processing of the Leucegene dataset was performed using 8 threads on a high-performance computing cluster running Centos 6.9.

#### Supplementary References


1. Eddelbuettel D, Francois R. Rcpp: Seamless R and C++ integration. Journal of Statistical Software. 2011;40(8):1 - 18.
2. Middleton R, Gao D, Thomas A, Singh B, Au A, Wong JJ, et al. IRFinder: assessing the impact of intron retention on mammalian gene expression. Genome Biol. 2017;18(1):51.
3. Ritchie ME, Phipson B, Wu D, Hu Y, Law CW, Shi W, et al. limma powers differential expression analyses for RNA-sequencing and microarray studies. Nucleic Acids Res. 2015;43(7):e47.
4. Love MI, Huber W, Anders S. Moderated estimation of fold change and dispersion for RNA-seq data with DESeq2. Genome Biol. 2014;15(12):550.
5. Ruddy S, Johnson M, Purdom E. Shrinkage of dispersion parameters in the binomial family, with application to differential exon skipping. The Annals of Applied Statistics. 2016;10(2):690-725.
6. Green ID, Pinello N, Song R, Lee Q, Halstead JM, Kwok CT, et al. Macrophage development and activation involve coordinated intron retention in key inflammatory regulators. Nucleic Acids Res. 2020;48(12):6513-29.
7. Patro R, Duggal G, Love MI, Irizarry RA, Kingsford C. Salmon provides fast and bias-aware quantification of transcript expression. Nat Methods. 2017;14(4):417-9.
8. Lavallée VP, Baccelli I, Krosl J, Wilhelm B, Barabé F, Gendron P, et al. The transcriptomic landscape and directed chemical interrogation of MLL-rearranged acute myeloid leukemias. Nat Genet. 2015;47(9):1030-7.
9. Dobin A, Davis CA, Schlesinger F, Drenkow J, Zaleski C, Jha S, et al. STAR: ultrafast universal RNA-seq aligner. Bioinformatics. 2013;29(1):15-21.
10. Liao Y, Smyth GK, Shi W. The R package Rsubread is easier, faster, cheaper and better for alignment and quantification of RNA sequencing reads. Nucleic Acids Research. 2019;47(8):e47-e.


#### Supplementary Figure Legends

**Figure S1: Forms of alternate splicing events** Included (Incl) and excluded (Excl) isoforms are defined as indicated.

**Figure S2: The SpliceWiz suite (A)** *SpliceWiz* consists of the *ompBAM* application programming interface (API*)*, command line interface (CLI), and the graphical user interface (GUI) as shown in this abbreviated package dependency map. **(B)** Reference generation using the *SpliceWiz* graphics user interface (GUI). Drop-down boxes allow interactive browsing of the Ensembl FTP site for the reference genome and gene annotation files. **(C)** Interactive experiment annotation using the SpliceWiz *GUI*. **(D)** Interactive volcano plot of differential alternative splicing events (ASEs). Users can select their differential ASE of interest using lasso or box select tool. These events will in turn be highlighted on the inter-connected scatter plots **(E)** and heatmaps **(F)**. Note the tooltip (triggered by mouse hover

over the heatmap cell) displays the sample name, splicing event name as well as the corresponding PSI value.

**Figure S3: Schematic of the COV file format.** Chromosome information is stored in the file header. Genomic coordinates and file offset pairs are stored in the index, allowing rapid recall of data contained in the file body, stored as discrete BGZF-compressed blocks. Only blocks containing the desired data are decompressed when data is read.

**Figure S4: Differences in ASE annotations between various tools (A)** Number of evaluable alternative splicing events (ASEs) in the percent-spliced-in (PSI) accuracy benchmark performed on the simulated dataset. **(B)** Number of ASEs derived from the transcript annotation by the evaluated tools. Alternate 5' / 3' splice site annotations of *rMATS* were matched with alternate first / last exon annotations of other tools if they contained matching splice junction pairs. **(C)** Overlap between ASE annotations of various tools. **(D)** Schematic illustrating the differences between annotation strategy of retained introns of *SpliceWiz* and *rMATS* / *SUPPA2*.

**Figure S5: Differential alternative splicing analysis of the Leucegene dataset using *SpliceWiz*** **(A)** Per-sample runtimes of each phase of the *SpliceWiz* pipeline. “Differential ASEs” refer to the per-sample runtime differential alternative splicing event (ASE) analysis using both *limma* and *DoubleExpSeq* function wrappers via *SpliceWiz*, for a single comparison between two experimental groups. Note that runtimes are expected to be roughly proportional to the sample size for all steps except differential ASE analysis. **(B)** *SRSF10* expression in the Leucegene dataset, divided by equal-sized expression bins (Low, Inter = Intermediate, High). **(C)** Differential ASE analysis between high and low *SRSF10*-expressing groups in the Leucegene dataset, using *limma* to model junction reads as log-normal distributed counts. A positive log2-fold change indicates upregulation of the *Included* isoform in high *SRSF10* samples. cpm = counts per million.

**Figure S6 Differential alternative splicing event analysis.** Differential alternative splicing events (ASEs) between high and low *SRSF10*-expressing groups in the Leucegene dataset, using *DoubleExpSeq* to model junction reads as beta-binomial distributed counts. A positive maximum likelihood expectation log2-fold change indicates upregulation of the *Included* isoform in high *SRSF10*-expressing samples. cpm = counts per million.

**Figure S7: Group-averaged coverage plots of differentially expressed alternative splicing events.** High *SRSF10* expression in the Leucegene dataset is associated with **(A)** increased *EXOSC10* intron retention, **(B)** increased skipping of *CTR9* alternate exon, and **(C)** increased usage of *FYN* upstream mutually exclusive exon. For each figure, the top panel illustrates coverage of two samples in each of high and low *SRSF10*-expressing groups; the middle panel shows group-averaged coverages of high, intermediate (inter) and low *SRSF10*-expressing groups; and the bottom panel shows stacked coverage traces of all 3 groups in the same plot.

**Figure S8: The *SpliceOver* parameter and its impact on intron retention quantitation. (A)** Theoretical aligned reads (top) and transcript annotation (bottom) flagged as included or excluded in *IRFinder's SpliceMax* or *SpliceWiz's SpliceOver* metrics. *SpliceOver* additionally accounts for spliced transcripts that overlap the intron of interest but share neither splice site. **(B)** Scatter plot comparing IR-ratios (of sample *01H002* from the Leucegene dataset) computed using *IRFinder's SpliceMax* and *SpliceWiz's SpliceOver* metrics. Red and black points represent IR events with discordant and concordant IR-ratios (ΔIR-ratio above and below 0.1) respectively. Note the events in the bottom right-hand side are over-estimated IR events that are corrected using SpliceWiz's *SpliceOver* metric.

#### Supplementary Tables

##### Supplementary Table 1: The *COV* format

| Field | Description | Type | Value |
|---|---|---|---|
| **COV Header**: contains the number, names and lengths of the chromosomes / scaffolds. Analogous to BAM format. BGZF compressed | | | |
| magic | COV file magic string (equivalent to BAM) | char[4] | COV\1 |
| n_ref | # reference sequences | uint32_t | $< 2^{31}$ |
| l_name | Length of the reference name plus 1 (including NUL) | uint32_t | |
| name | Reference sequence name; NUL-terminated | char[l_name] | |
| l_ref | Length of the reference sequence, in nucleotides | uint32_t | |
| **COV Index**: recording chromosome and start coordinate of each BGZF block compressed in the body. BGZF compressed. Positive strand is recorded first, followed by negative followed by unstranded, thus giving 3 x n_ref number of index blocks | | | |
| index_size | Total length of the index entry for each chromosome, excluding this field | uint32_t | |
| block_start | The first genomic coordinate of the BGZF block | uint32_t | |
| file_offset | File offset in of the compressed BGZF block, in bytes, from the beginning of the index block | uint64_t | |
| **COV Body**: Contains coverage values and lengths in RLE format. The start coordinates of the first value of every uncompressed block is given by the COV index. | | | |
| BGZF block | | | |
| cov_value | Depth of RNA-seq coverage | int32_t | |
| cov_length | Number of nucleotides with the above coverage value. The start coordinate of this is the end coordinate of the last record. If this is the first record, the start coordinate is given by the block_start value in the index. | uint32_t | |
| **BGZF end-of-file terminator** | | | |

**Supplementary Table 2A: SpliceWiz Filter Parameters**

| Filter | Setting | Parameter (non-IR) | Parameter (IR) | Purpose |
|---|---|---|---|---|
| Depth | - | 20 | 20 | Removes all ASEs with transcript abundance (sum of included and excluded isoforms) is less than this amount. |
| Participation | Soft | 0.4 | 0.7 | For IR, participation refers to the proportion of measurable intron covered by RNA-seq reads. The filter is applied to samples if the mean intronic abundance is at least 5.<br>For other ASEs, it refers to the proportion of total transcript depth of spliced transcripts that is accounted to either the inclusion or exclusion isoform.<br>The filter is applied to all ASEs with a transcript abundance (included + excluded reads) of at least 20. |
| | Medium | 0.6 | 0.8 | |
| | Hard | 0.8 | 0.9 | |
| Consistency | Soft | 0.25 | 0.25 | Consistency refers to the comparative orders of magnitude of the two tandem junctions.<br>In IR, these are the flanking exon-intron overhang reads.<br>Tandem junctions / overhang reads belonging to an isoform are deemed consistent if the consistency ratio is at larger than or equal to this parameter.<br>The filter is applied to all ASEs with a transcript abundance of at least 20. |
| | Hard | 0.5 | 0.5 | |

**Supplementary Table 2B: Optimized SpliceWiz filters**

| Filter | Setting | Parameter (non-IR) | Parameter (IR) | Applicable ASEs |
|---|---|---|---|---|
| Depth | - | 20 | 20 | All |
| Participation | Soft | - | 0.7 | SE, A5SS, A3SS |
| Consistency | Soft | 0.25 | 0.25 | IR, SE, MXE |
| Terminal Exon | - | - | - | AFE, ALE |

**Supplementary Table 3: Parameters used for flux-simulator**

| Parameter | Value |
|---|---|
| TSS_MEAN | 25 |
| POLYA_SCALE | NaN |
| POLYA_SHAPE | NaN |
| FRAG_SUBSTRATE | RNA |
| FRAG_METHOD | UR |
| FRAG_UR_ETA | NaN |
| FRAG_UR_DELTA | NaN |
| FRAG_UR_D0 | 3 |
| RTRANSCRIPTION | YES |
| RT_MOTIF | Default |
| RT_PRIMER | RH |
| RT_LOSSLESS | YES |
| RT_MIN | 500 |
| RT_MAX | 5500 |
| FILTERING | YES |
| PCR_PROBABILITY | 0.05 |
| GC_MEAN | NaN |
| GC_SD | NaN |
| READ_NUMBER | 200,000,000 |
| READ_LENGTH | 100 |
| PAIRED_END | YES |
| FASTA | YES |
| ERR_FILE | 76 |

**Supplementary Table 4: Commands and parameters used for BAM processing benchmarks**

| Tool | Command / Parameters |
|---|---|
| rMATS | `python rmats.py -t paired --readLength 50 \`<br>` --variable-read-length --allow-clipping \`<br>` --nthread 4 --task prep` |
| MAJIQ | `majiq build -j 4 -c config.txt`<br><br>(inside config.txt)<br>`genome=hg38`<br>`readlen=100`<br>`strandedness=reverse` |
| IRFinder | Below run inside `BiocParallel::bplapply()` with `MulticoreParam(2)` (as each instance of IRFinder consumes two threads):<br><br>`gzip -cd < ${bamFile} \| \`<br>` bin/util/irfinder {defaults for IRFinder reference}` |
| IntEREst | `interest(`<br>` isPaired = TRUE, method = c("IntRet", "IntSpan"),`<br>` bpparam = MulticoreParam(4)`<br>`)` |
| ASpli | `gbc <- gbCounts(`<br>` minReadLength = 100, maxISize = 50000,`<br>` libType = "PE", strandMode = 2`<br>`)`<br>`jCounts(`<br>` gbc, minReadLength = 100,`<br>` libType = "PE", strandMode = 2`<br>`)` |
| SGSeq | `sgfc <- analyzeFeatures(…, cores = 4)`<br>`sgv <- analyzeVariants(sgfc, cores = 4)`<br>`sgv_ranges <- rowRanges(sgv)`<br>`sgvc <- getSGVariantCounts(sgv_ranges, cores = 4)` |

### Figure S1

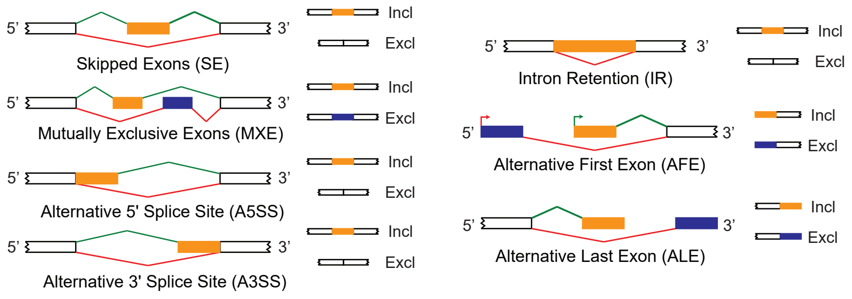

5'
3'
Incl
Excl
Skipped Exons (SE)
Mutually Exclusive Exons (MXE)
Alternative 5' Splice Site (A5SS)
Alternative 3' Splice Site (A3SS)
Intron Retention (IR)
Alternative First Exon (AFE)
Alternative Last Exon (ALE)

Figure S2

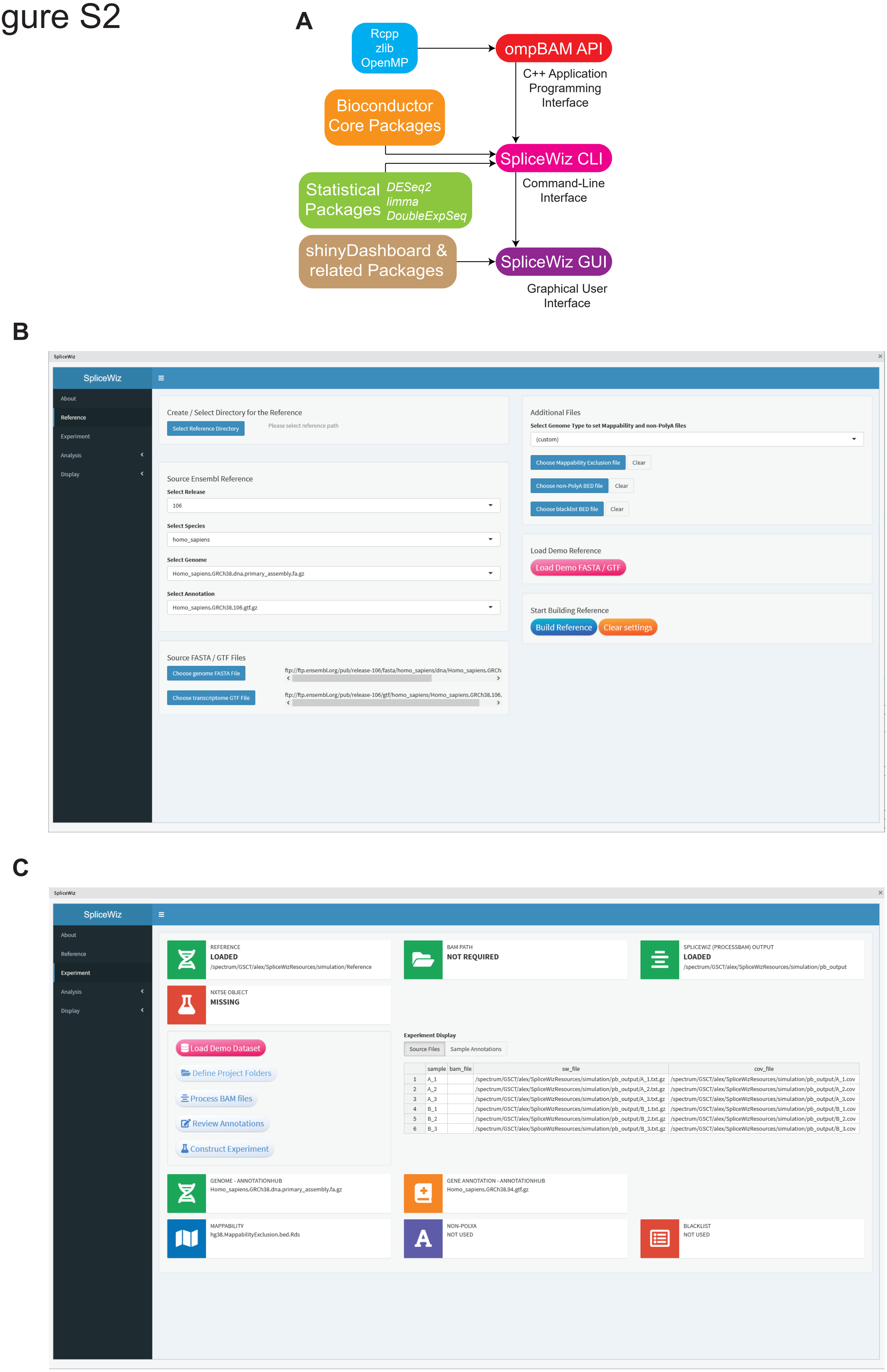
A
Rcpp
zlib
OpenMP
ompBAM API
C++ Application
Programming
Interface
Bioconductor
Core Packages
SpliceWiz CLI
Statistical Packages
DESeq2
limma
DoubleExpSeq
Command-Line
Interface
shinyDashboard &
related Packages
SpliceWiz GUI
Graphical User
Interface
B
SpliceWiz
About
Reference
Experiment
Analysis
Display
Create / Select Directory for the Reference
Select Reference Directory
Please select reference path
Source Ensembl Reference
Select Release
106
Select Species
homo_sapiens
Select Genome
Homo_sapiens.GRCh38.dna.primary_assembly.fa.gz
Select Annotation
Homo_sapiens.GRCh38.106.gtf.gz
Source FASTA / GTF Files
Choose genome FASTA File
Choose transcriptome GTF File
Additional Files
Select Genome Type to set Mappability and non-PolyA files
(custom)
Choose Mappability Exclusion file
Clear
Choose non-PolyA BED file
Choose blacklist BED file
Load Demo Reference
Load Demo FASTA / GTF
Start Building Reference
Build Reference
Clear settings
C
SpliceWiz
REFERENCE
LOADED
BAM PATH
NOT REQUIRED
SPLICEWIZ (PROCESSBAM) OUTPUT
LOADED
NXTSE OBJECT
MISSING
Load Demo Dataset
Define Project Folders
Process BAM files
Review Annotations
Construct Experiment
Experiment Display
Source Files
Sample Annotations
GENOME - ANNOTATIONHUB
Homo_sapiens.GRCh38.dna.primary_assembly.fa.gz
GENE ANNOTATION - ANNOTATIONHUB
Homo_sapiens.GRCh38.94.gtf.gz
MAPPABILITY
hg38.MappabilityExclusion.bed.Rds
NON-POLYA
NOT USED
BLACKLIST
NOT USED

### Figure S2

**D**

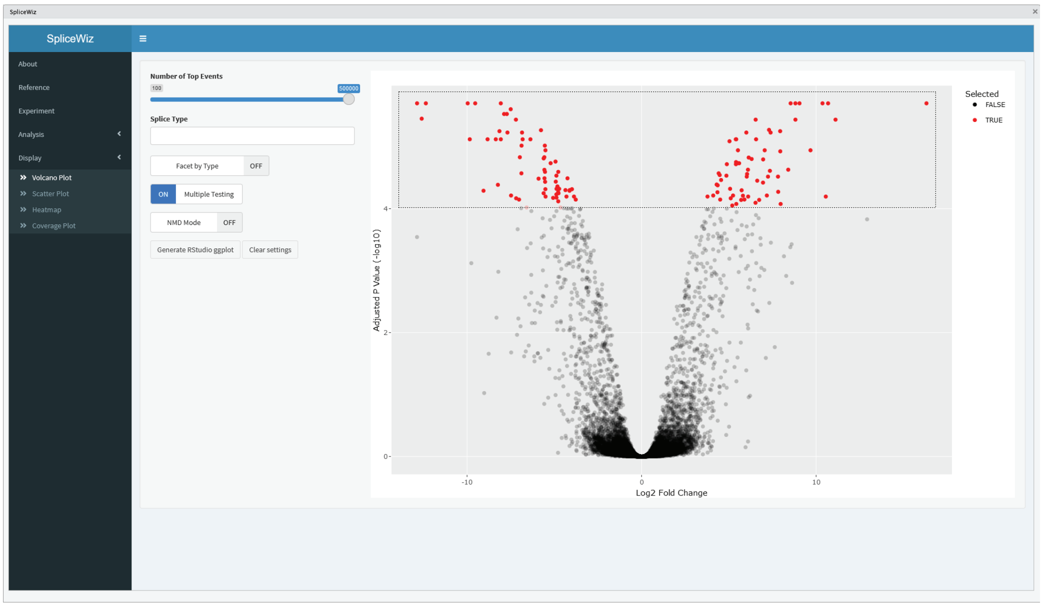


**E**

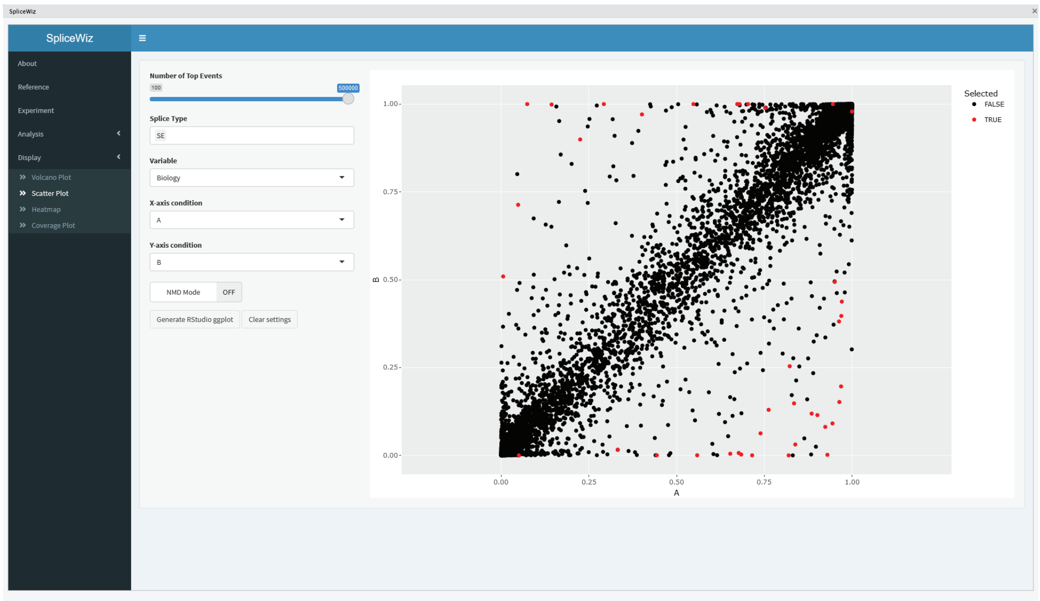


**F**

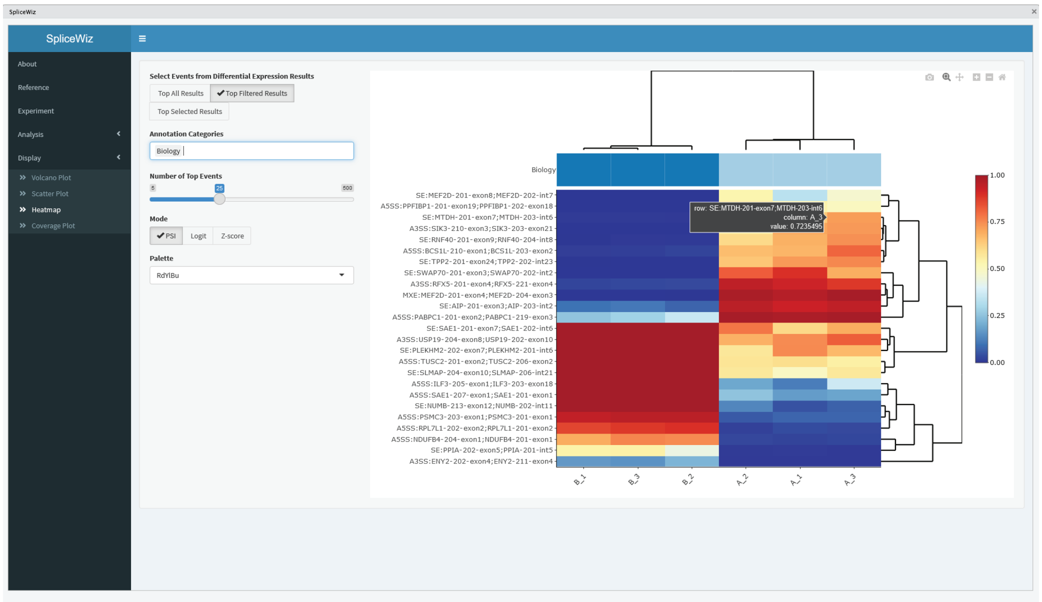

### Figure S3

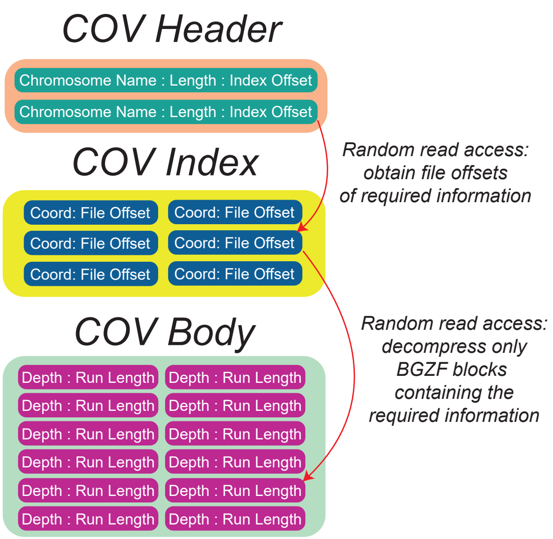
COV Header
Chromosome Name : Length : Index Offset
Chromosome Name : Length : Index Offset
COV Index
Coord: File Offset
COV Body
Depth : Run Length
Random read access: obtain file offsets of required information
Random read access: decompress only BGZF blocks containing the required information

### Figure S4

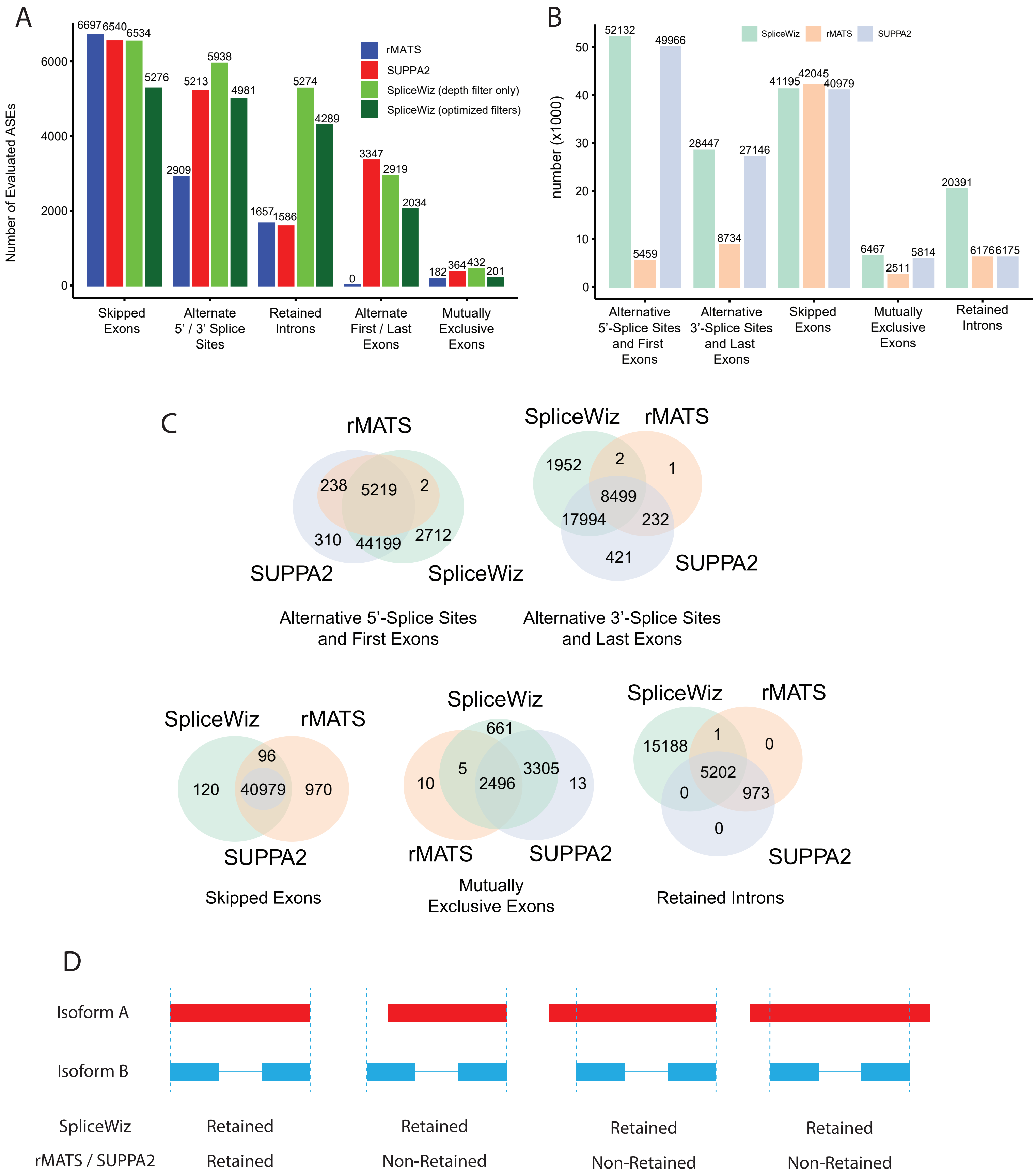
A
Number of Evaluated ASEs
0
2000
4000
6000
rMATS
SUPPA2
SpliceWiz (depth filter only)
SpliceWiz (optimized filters)
6697
6540
6534
5276
2909
5213
5938
4981
1657
1586
5274
4289
0
3347
2919
2034
182
364
432
201
Skipped Exons
Alternate 5' / 3' Splice Sites
Retained Introns
Alternate First / Last Exons
Mutually Exclusive Exons
B
number (x1000)
0
10
20
30
40
50
SpliceWiz
rMATS
SUPPA2
52132
5459
49966
28447
8734
27146
41195
42045
40979
6467
2511
5814
20391
6176
6175
Alternative 5'-Splice Sites and First Exons
Alternative 3'-Splice Sites and Last Exons
Skipped Exons
Mutually Exclusive Exons
Retained Introns
C
rMATS
238
5219
2
310
44199
2712
SUPPA2
SpliceWiz
Alternative 5'-Splice Sites and First Exons
SpliceWiz
rMATS
1952
2
1
8499
17994
232
421
SUPPA2
Alternative 3'-Splice Sites and Last Exons
SpliceWiz
rMATS
96
120
40979
970
SUPPA2
Skipped Exons
SpliceWiz
661
5
3305
10
2496
13
rMATS
SUPPA2
Mutually Exclusive Exons
SpliceWiz
rMATS
15188
1
0
5202
0
973
0
SUPPA2
Retained Introns
D
Isoform A
Isoform B
SpliceWiz
rMATS / SUPPA2
Retained
Retained
Retained
Non-Retained
Retained
Non-Retained
Retained
Non-Retained

### Figure S5

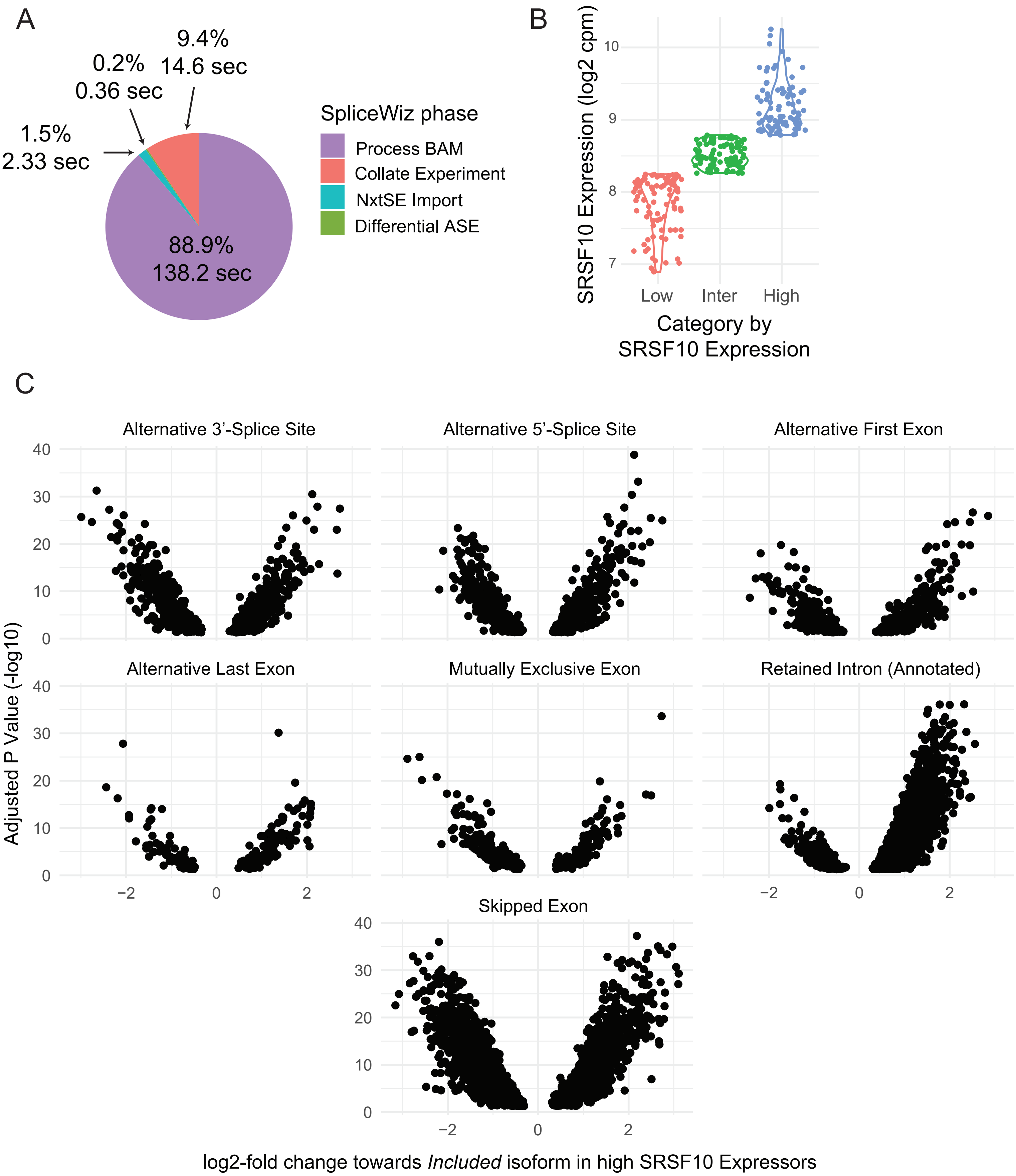
A
9.4%
14.6 sec
0.2%
0.36 sec
1.5%
2.33 sec
88.9%
138.2 sec
SpliceWiz phase
Process BAM
Collate Experiment
NxtSE Import
Differential ASE
B
SRSF10 Expression (log2 cpm)
Low
Inter
High
Category by
SRSF10 Expression
C
Alternative 3'-Splice Site
Alternative 5'-Splice Site
Alternative First Exon
Alternative Last Exon
Mutually Exclusive Exon
Retained Intron (Annotated)
Skipped Exon
Adjusted P Value (-log10)
log2-fold change towards *Included* isoform in high SRSF10 Expressors

### Figure S6

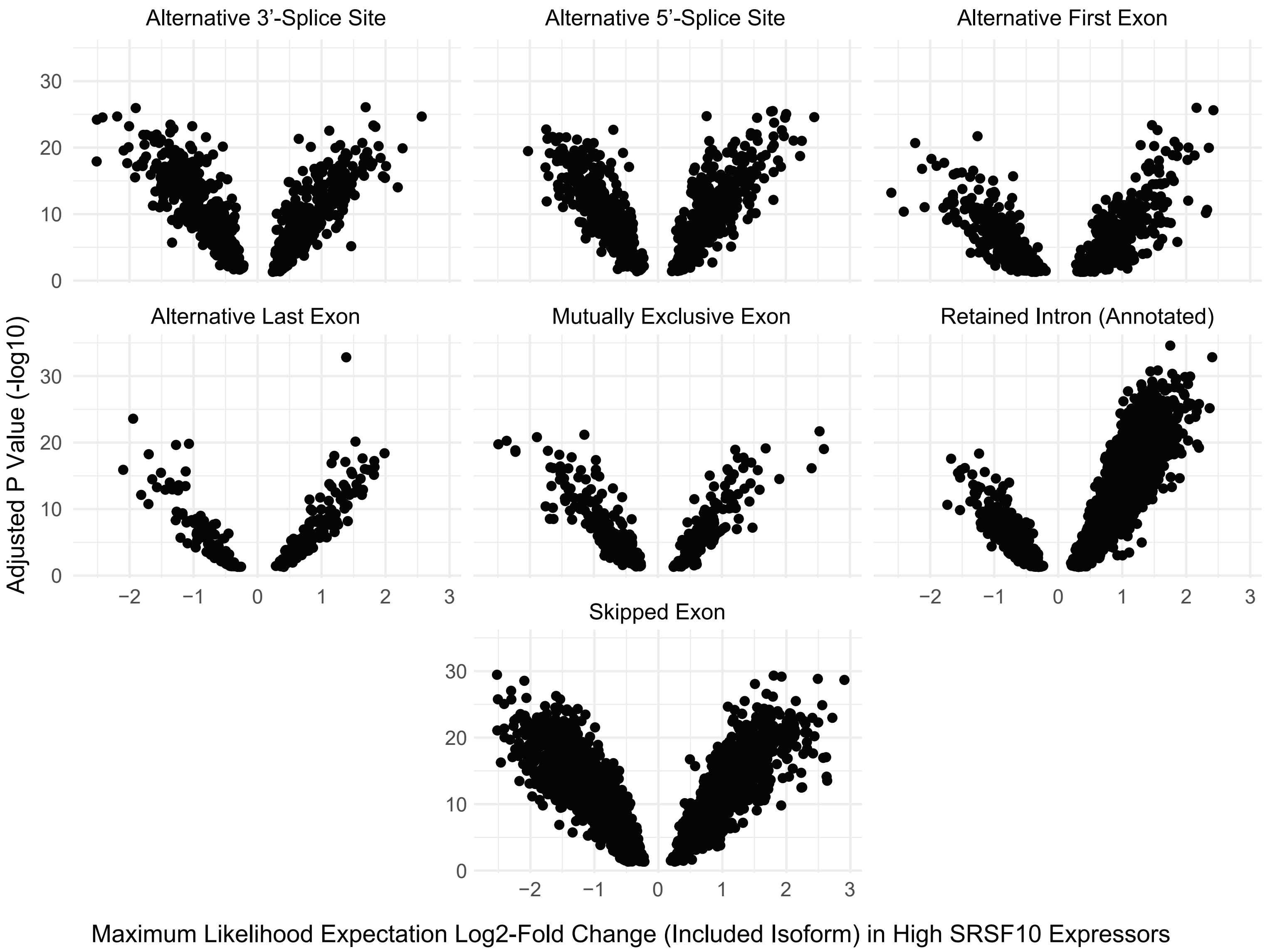
Alternative 3'-Splice Site
Alternative 5'-Splice Site
Alternative First Exon
Alternative Last Exon
Mutually Exclusive Exon
Retained Intron (Annotated)
Skipped Exon
Adjusted P Value (-log10)
Maximum Likelihood Expectation Log2-Fold Change (Included Isoform) in High SRSF10 Expressors

### Figure S7

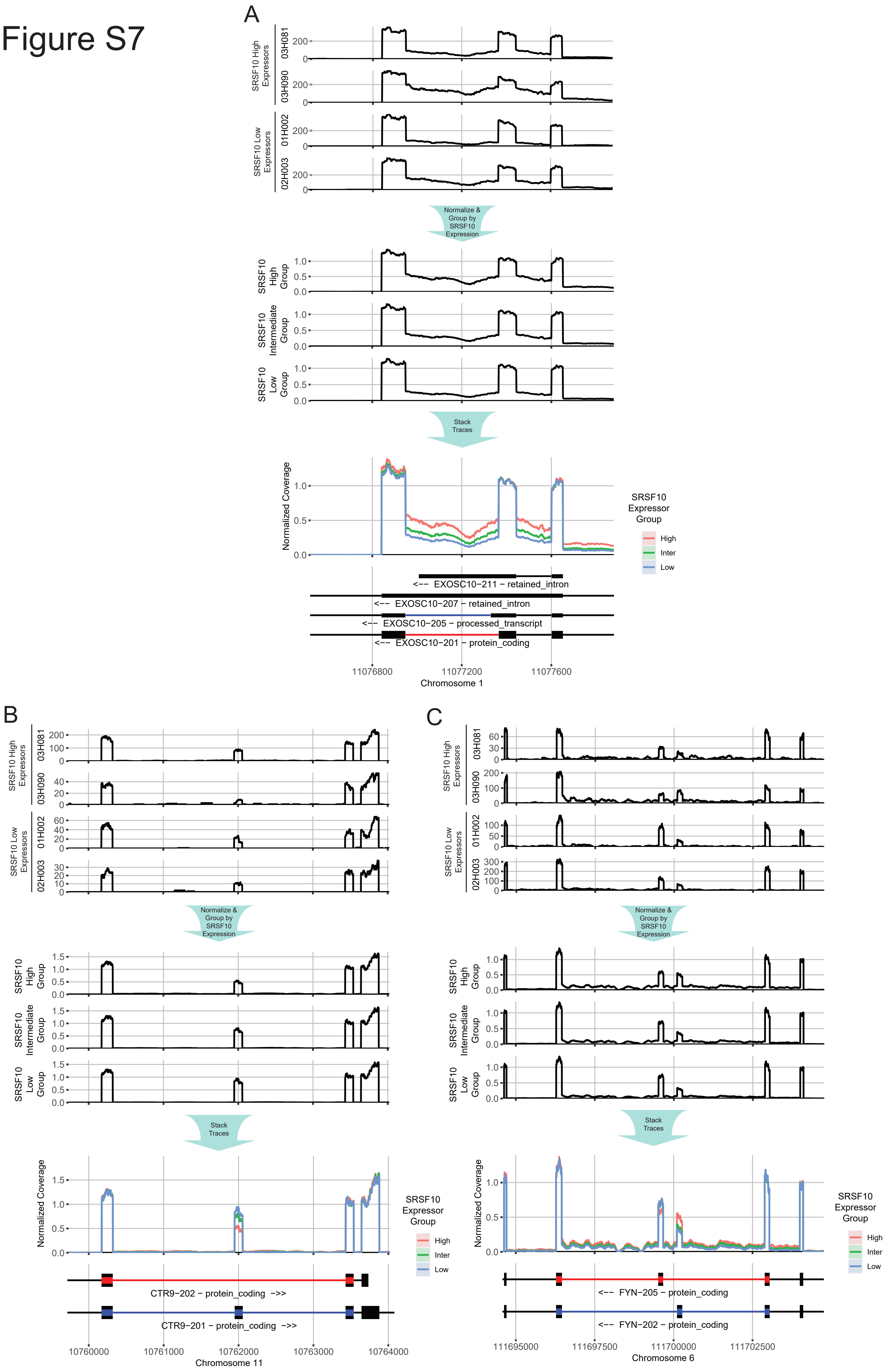

A
SRSF10 High Expressors
03H081
03H090
SRSF10 Low Expressors
01H002
02H003
Normalize & Group by SRSF10 Expression
SRSF10 High Group
SRSF10 Intermediate Group
SRSF10 Low Group
Stack Traces
Normalized Coverage
SRSF10 Expressor Group
High
Inter
Low
<-- EXOSC10-211 - retained_intron
<-- EXOSC10-207 - retained_intron
<-- EXOSC10-205 - processed_transcript
<-- EXOSC10-201 - protein_coding
11076800
11077200
11077600
Chromosome 1
B
CTR9-202 - protein_coding ->>
CTR9-201 - protein_coding ->>
10760000
10761000
10762000
10763000
10764000
Chromosome 11
C
<-- FYN-205 - protein_coding
<-- FYN-202 - protein_coding
111695000
111697500
111700000
111702500
Chromosome 6

### Figure S8

A

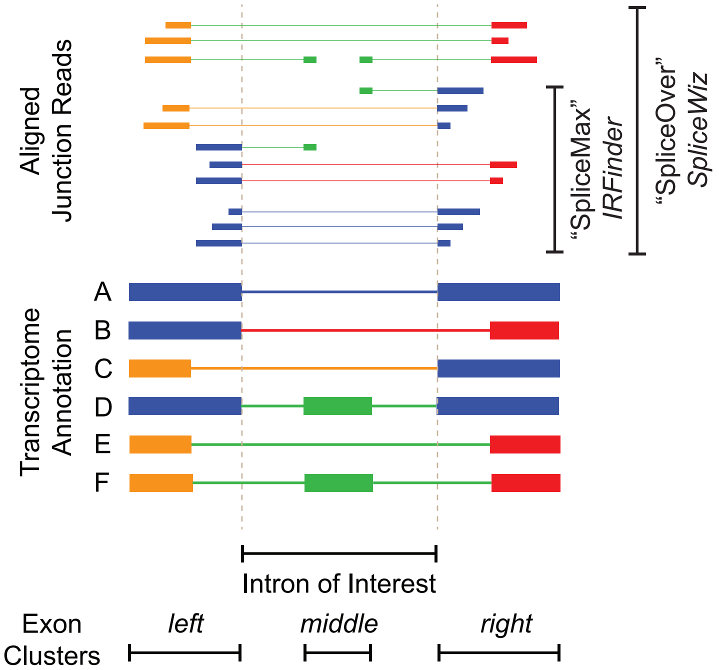
Aligned Junction Reads
"SpliceMax"
IRFinder
"SpliceOver"
SpliceWiz
Transcriptome Annotation
A
B
C
D
E
F
Intron of Interest
Exon Clusters
left
middle
right

B

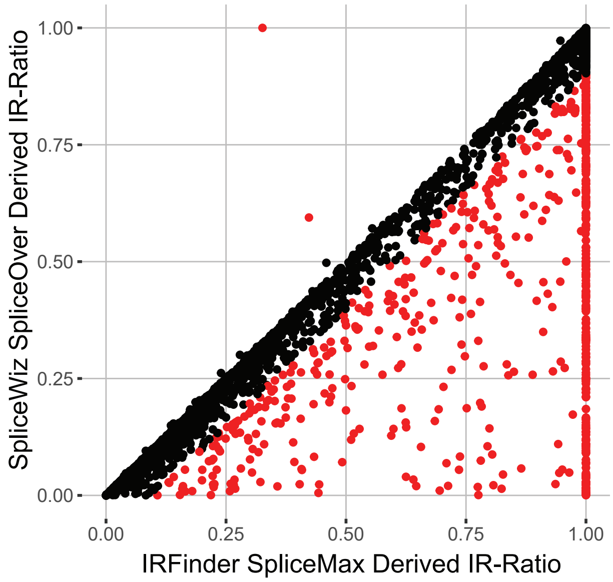
SpliceWiz SpliceOver Derived IR-Ratio
1.00
0.75
0.50
0.25
0.00
0.00
0.25
0.50
0.75
1.00
IRFinder SpliceMax Derived IR-Ratio
